# Detection and characterization of apoptotic and necrotic cell death by time-lapse quantitative phase image analysis

**DOI:** 10.1101/589697

**Authors:** Tomas Vicar, Martina Raudenska, Jaromir Gumulec, Michal Masarik, Jan Balvan

**Affiliations:** Department of Physiology, Faculty of Medicine, Masaryk University / Kamenice 5, CZ-625 00 Brno, Czech Republic; Department of Pathological Physiology, Faculty of Medicine, Masaryk University / Kamenice 5, CZ-625 00 Brno, Czech Republic; Central European Institute of Technology, Brno University of Technology, Technicka 3058/10, CZ-616 00 Brno, Czech Republic; Department of Biomedical Engineering, Faculty of Electrical Engineering and Communication, Brno University of Technology, Technicka 3058/10, Brno, Czech Republic; BIOCEV, First Faculty of Medicine, Charles University, Průmyslová 595, 252 50, Vestec, Czech Republic

**Keywords:** cell death, apoptosis, necrosis, Coherence-Controlled Holographic Microscope, Quantitative Phase Imaging, cell dry mass, cell morphology, correlative microscopy, machine learning, Staurosporine, Doxorubicin, black phosphorus

## Abstract

Cell viability and cytotoxicity assays are highly important for drug screening and cytotoxicity tests of antineoplastic or other therapeutic drugs. Even though biochemical-based tests are very helpful to obtain preliminary preview, their results should be confirmed by methods based on direct cell death assessment. In this study, time-dependent changes in quantitative phase-based parameters during cell death were determined and methodology useable for rapid and label-free assessment of direct cell death was introduced. Our method utilizes Quantitative Phase Imaging (QPI) which enables the time-lapse observation of subtle changes in cell mass distribution. According to our results, morphological and dynamical features extracted from QPI micrographs are suitable for cell death detection (76% accuracy in comparison with manual annotation). Furthermore, based on QPI data alone and machine learning, we were able to classify typical dynamical changes of cell morphology during both caspase 3,7-dependent and independent cell death subroutines. The main parameters used for label-free detection of these cell death modalities were cell density (pg/pixel) and average intensity change of cell pixels further designated as Cell Dynamic Score (CDS). To the best of our knowledge, this is the first study introducing CDS and cell density as a parameter typical for individual cell death subroutines with prediction accuracy 75.4 % for caspase 3,7-dependent and -independent cell death.

## Introduction

Analysis of cell viability and the distinction of specific cell death subtype represents a key aspect in many areas of cell biology. This kind of information is also highly important for drug screening and cytotoxicity tests of antineoplastic or other therapeutic drugs. Cell death is considered reversible until a first ‘point-of-no-return’ is overstepped. While no exactly defined biochemical event can be taken as an undisputable proof of this point-of-no-return, cell should be taken as dead when any of these situations occur: (1) the cell has lost the integrity of its plasma membrane; (2) the cell and its nucleus has undergone complete fragmentation into discrete bodies (apoptotic bodies); (3) the cellular corpse has been engulfed and digested by a neighbouring cell ^1^. Cells that are arrested in the cell cycle should be counted as viable ^2^.

Methods for cell death analysis are usually based on various basic cell functions such as enzyme activity, semi-permeability of the mitochondrial or cellular membrane, cell adherence, ATP production, the presence of specific markers, or changes of functionality due to specific inhibitor (genetical or pharmacological) ^3^. Methods for cell death detection form two main groups: a) methods that directly measure cell death; and b) methods that analyse biochemical processes or features characteristic for viable cells ^4^. Even though indirect tests are very helpful to obtain preliminary preview, their results should be confirmed by methods based on direct cell death assessment ^3^. According to NCCD (Nomenclature Committee on Cell Death), the currently accepted definition of cell death and its subroutines is based particularly on genetical, biochemical, pharmacological, and functional parameters, rather than morphological aspects ^1^. Specific methods for apoptosis and necrosis detection are focused on typical biochemical parameters such as visualization of phosphatidylserine exposure, executioner caspases activation, or DNA fragmentation in the case of apoptotic cell death; and loss of barrier function and subsequent permeabilization of the plasma membrane with the release of specific death associated molecular patterns (DAMP) during necrotic cellular demise ^1^. Nevertheless, almost all these methods are based on fluorometric or colourimetric endpoint visualization of the analyzed parameter and belong among indirect assays. Such indirect endpoint analyses are prone to the misleading results as we have shown in our previous work ^5^. Although morphological aspects of cell death are not generally recommended to determine cell death subroutines ^1^, it would be a mistake to completely ignore them. Recent progress in Quantitative Phase Imaging techniques (QPI) has enabled the observation of time-dependent subtle changes unrecognizable to the naked eye (such as cell mass distribution) on micrographs. These changes in cell mass distribution, cell density, micro-blebbing of the cell membrane, nuclear shape, homogeneity of cell content, and many other parameters, which can be typical for individual cell death subroutines, can be observed without fixation, labelling or cell harvesting.

In this article, we demonstrate a methodology useable for rapid assessment of direct cell death that is based on NCCD recommendations. Using advanced quantitative label-free phase imaging (QPI), we were also able to observe typical dynamical changes of cell morphology during both caspase 3, 7-dependent and -independent cell death subroutines and to determine the moment of cell death. Dynamical, time-dependent changes in cell mass distribution maps were used for the label-free distinction between typical morphological patterns appropriate for caspase-dependent and caspase-independent cell death subroutines.

As model cells, prostate cancer cell lines DU145, LNCaP, and benign cell line PNT1A established by immortalization of normal adult prostatic epithelial cells were used. These cell lines were not selected because of the clinical relevance, but rather because of their radically different size and accessibility for the automatic segmentation algorithm. We expected cellular size to be the factor influencing the rate of morphological changes during cell death. Cell death was triggered by doxorubicin and staurosporine, two commonly known inducers of apoptosis with a distinct induction of caspase cleavage and consequently a distinct morphological manifestation ^6, 7^. Cell death induced by black phosphorus (BP) was detected as an example of difficult detection conditions ^8^.

Cell tracking is an essential step in cell image analysis. Even though many methods exist ^9^, there is no universal and sufficiently robust method applicable to QPI, especially for touching cells and if correct tracking through whole image sequence is needed. Simple available tools like TrackMate ^10^ have shown to be insufficient, thus we have developed a new tracking method tailored for our dataset.

The number of existing methods for the detection of cell death in label-free time-lapse images is very limited. In ^11^ authors detect cell death event in Phase Contrast Microscopy using features time-series (size, roundness, speed etc.) and classify each time point as alive or death with transductive support vector machine. In comparison, our technique based on quantitative microscopy data uses more features and more advanced Long Short-Term Memory (LSTM) neural network.

There are several techniques for static cell image classification including extraction of features and application of classifier ^12, 13^ or application of convolutional neural network ^14^. These techniques can be also possibly applied for a distinction of different types of cell death, however, we decided for using only two features for this distinction in order to make our results easily interpretable, while also incorporating new features describing cell dynamics.

## Methods

### Chemical and biochemical reagents

RPMI-1640 medium, fetal bovine serum (FBS) (mycoplasma-free), penicillin/streptomycin, and trypsin were purchased from Sigma Aldrich Co. (St. Louis, MO, USA). Phosphate buffered saline PBS was purchased from Invitrogen Corp. (Carlsbad, CA, USA). Ethylenediaminetetraacetic acid (EDTA), staurosporine, doxorubicin and all other chemicals of ACS purity were purchased from Sigma Aldrich Co. (St. Louis, MO, USA) unless noted otherwise. Black phosphorus was kindly provided by Dr. Martin Pumera.

### Cell culture and cultured cell conditions

LNCaP cell line was established from a lymph node metastase of the hormone-refractory patient and contains a mutation in the androgen receptor (AR) gene. This mutation creates a promiscuous AR that can bind to different types of steroids. LNCaP cells are AR-positive, PSA-positive, PTEN-negative and harbor wild-type p53 ^15, 16^. PNT1A is an immortalized non-tumorigenic epithelial cell line. PNT1A cells harbour wild-type p53. However, SV40 induced T-antigen expression inhibits the activity of p53. This cell line had lost the expression of AR and prostate-specific antigen (PSA) ^17^. DU-145 cell line is derived from the metastatic site in the brain and contains P223L and V274F mutations in p53. This cell line is PSA and AR-negative and androgen independent ^18^. All cell lines used in this study were purchased from HPA Culture Collections (Salisbury, UK) and were cultured in RPMI-1640 medium with 10 % FBS. The medium was supplemented with antibiotics (penicillin 100 U/ml and streptomycin 0.1 mg/ml). Cells were maintained at 37°C in a humidified (60%) incubator with 5% CO2 (Sanyo, Japan). Cell lines were not tested on mycoplasma contamination.

### Correlative time-lapse quantitative phase-fluorescence imaging

QPI and fluorescence imaging were performed by using multimodal holographic microscope Q-PHASE (TESCAN, Brno, Czech Republic). To determine the amount of caspase-3/7 product accumulation, cells were loaded with 2 µM CellEvent™ Caspase-3/7 Green Detection Reagent (Life Technologies, Carlsbad, CA, USA) according to the manufacturer’s protocol and visualized using FITC 488 nm filter. To detect the cells with a loss of plasma membrane integrity, cells were stained with 1 ug/ml propidium iodide (Sigma Aldrich Co., St. Louis, MO, USA) and visualized using TRITC 542 nm filter. Nuclear morphology and chromatin condensation were analyzed using Hoechst 33342 nuclear staining (ENZO, Lausen, Switzerland) and visualized using DAPI 461 nm filter. Cells were cultivated in Flow chambers μ-Slide I Lauer Family (Ibidi, Martinsried, Germany). To maintain standard cultivation conditions (37°C, humidified air (60%) with 5% CO2) during time-lapse experiments, cells were placed in the gas chamber H201 - for Mad City Labs Z100/Z500 piezo Z-stages (Okolab, Ottaviano NA, Italy). To image enough cells in one field of view, Nikon Plan 10/0.30 was chosen. For each of three cell lines and each of three treatments, seven fields of view were observed with the frame rate 3 mins/frame for 24 or 48 h respectively.

Holograms were captured by CCD camera (XIMEA MR4021 MC-VELETA), fluorescence images were captured using ANDOR Zyla 5.5 sCMOS camera. Complete quantitative phase image reconstruction and image processing were performed in Q-PHASE control software. Cell dry mass values were derived according to ^19^ and ^20^ from the phase (eq. (1)), where m is cell dry mass density (in pg/μm2), φ is detected phase (in rad), λ is wavelength in μm (0.65 μm in Q-PHASE), and α is specific refraction increment (≈0.18 μm^3^/pg). All values in the formula except the φ are constant. The value of φ (Phase) is measured directly by the microscope.

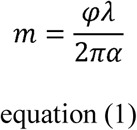

Integrated phase shift through a cell is proportional to its dry mass, which enables studying changes in cell mass distribution ^20^.

### Cell dry mass tracking

The custom method for automatic cell tracking and measuring of selected features was developed and implemented in MATLAB. Although tracking is not the main contribution of this article, it is a necessary step for our analysis and there is no available method suitable for QPI. The proposed tracking method consists of two main steps - nuclei tracking followed by expansion of each nucleus region to the whole cell.

Nuclei tracking is done by Gaussian Mixture Model (GMM) fitting to nuclei image (Hoechst 33342) in each frame by Expectation-Maximization (EM) algorithm (a similar method is used for segmentation in ^21^). For each frame, Gaussians are provided from the previous frame and their parameters are updated by several steps of EM. Before the application of GMM, background pixels are eliminated by segmentation using Maximally Stable Extremal Region (MSER) ^22^. Gaussians parameters are optimized for each nuclei cluster separately. In the first frame, positions of Gaussians are initialized manually and the covariance matrices are initialized as an identity matrix. The whole algorithm is summarized in Fig. 1 and for more details see ^23^. This method does not recognise division of cells. For the next analysis, we use only cells, which occur in the whole image sequence without division or joining with other cell (which appears mainly due to tracking and segmentation errors).

**Figure 1.**
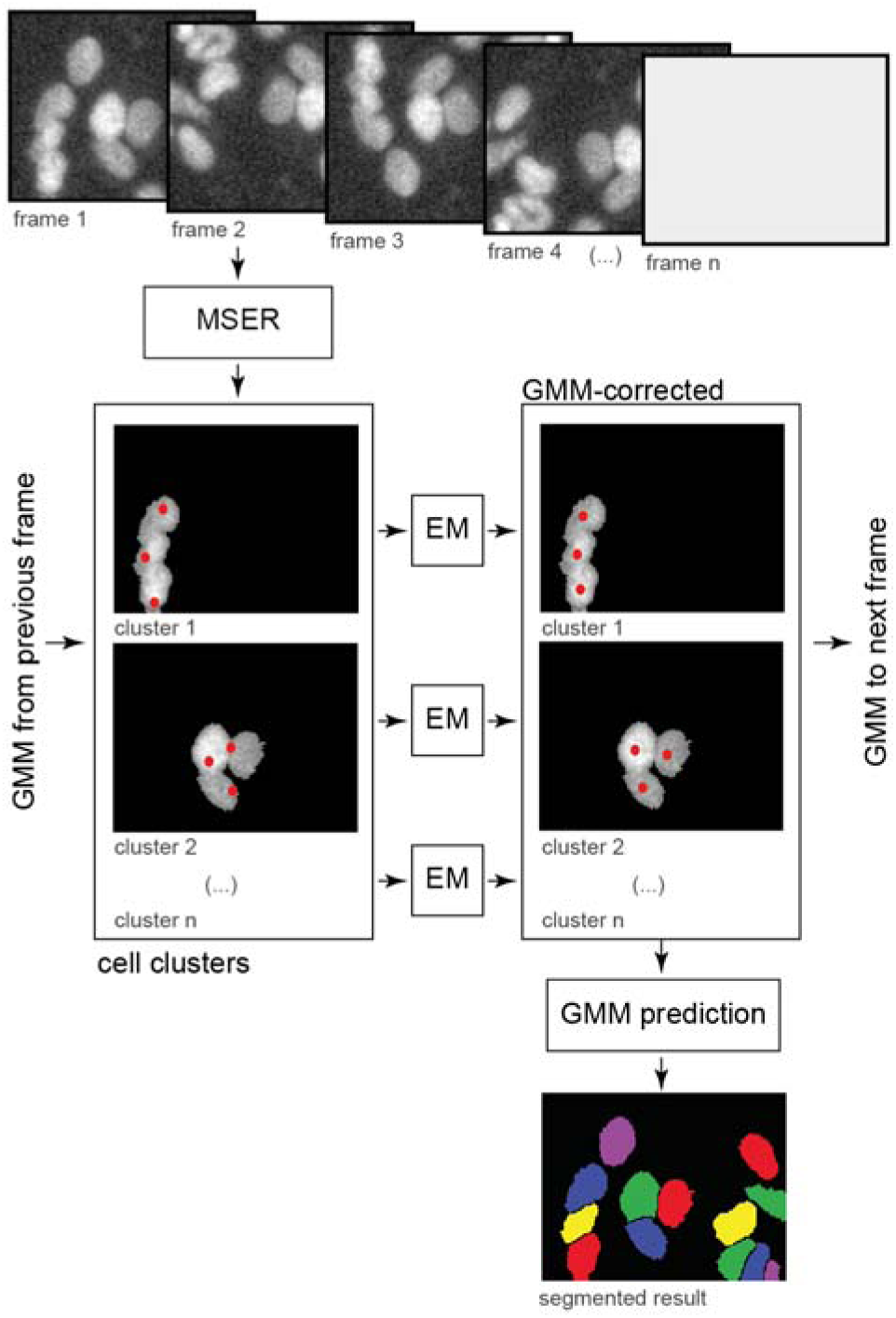
block diagram of cell tracking algorithm. Nuclear staining with Hoechst 33342 used as an input. EM, Expectation-Maximization; GMM, Gaussian Mixture Model; MSER, maximally stable extremal region;

The previous step provides tracked nuclei, but for extraction of most of the cell features, we need cell segmentation. However, nuclei can be used as seeds for segmentation in each frame. We can use simple thresholding for segmentation of foreground in QPI cell image, where the threshold value can be the same in all frames, which provides sufficient results for a distinction between foreground and background, but a separation of single cells is very problematic. We use tracked nuclei as seeds for proper division of the foreground binary mask *F*_*t*_ (obtained by tresholding) to the single cells binary mask *S*_*t*_. Seeded watershed ^24^ is used on the negative of the original image *O*_*t*_ with nuclei tracking result as seedpoints. Watershed results (boundary lines) are then used for division of the foreground mask.

However, the simple application of the watershed algorithm leads to a fast mass exchange between cells due to segmentation errors, which is very undesirable for precise feature extraction. For this reason, we introduced a simple modification named Movement Regularized Watershed (MRW), in order not to allow dramatic contour changes between frame. This can be achieved by incorporating the mask from the previous frame to the actual frames watershed calculation. This can be done by modifying *O*_*t*_ and *F*_*t*_ as

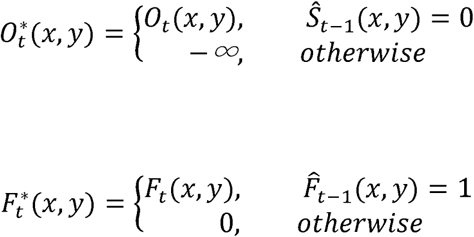

Where *Ŝ*_*t*-1_ (*x, y*) and 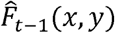 are an eroded versions of single cell segmentation and dilated version of foreground segmentation from the previous frame, respectively. Modified 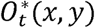 and 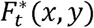 are used in the seeded watershed algorithm. Modification of 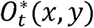 forbid high area exchange between cells in consecutive frames and modification of 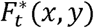 forbid high contour movement into the background between frames. Maximal possible contour movement than can be set by the amount of erosion and dilatation.

### Manual dataset annotation

For each cell line and each treatment, seven FOVs were processed by the tracking method and only complete tracklets were kept for manual annotation. Overall, 819 PNT1A, 755 DU-145, and 581 LNCaP cells with annotated cell death were analysed. Timepoints of cell death and apoptotic or necrotic morphotype were manually annotated by a skilled professional (JB). Following parameters were considered: Casp 3,7 signal, PI signal, nuclear morphology, plasma membrane rupture and blebbing, surface detachment and cell rounding. A total number of 230, 196 and 220 apoptotic morphotypes for DU-145, LNCaP and PNT1A, respectively, was detected. A total number of 421, 237 and 441 necrotic morphotypes for DU-145, LNCaP and PNT1A, respectively, was detected. Remaining cells survived the treatment.

### Feature extraction

For further analysis, we extracted several cell features including cell mass, area, mass density (average pixel brightness), cell speed (centroid movement), circularity, eccentricity and maximum of the histogram, the position of maxima of histogram and entropy of histogram. Besides the classical cell features, we introduce tailored feature Cell Dynamic Score (CDS). CDS is a mean Euclidian distance between cell pixels in the actual and the following frame computed as

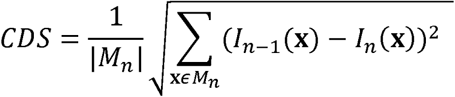

where *M*_*n*_ is a set of poisitons defined by the cell segmentation mask in the *n*-th frame and *I*_*n*_ is the *n*-th frame of QPI. CDS provides information about the speed of change of the cell pixel values due to both movement and morphological changes, but it is not much dependent on the segmentation quality, because the same mask is used in both frames. All these features were evaluated in all frames, where the result is a set of signals describing the cell behaviour in time.

### Label-free algorithm for cell death detection

Besides the significant mass decrease of dead cells, various feature evolvement types were observed during detected cell deaths, which complicate the expert specification of the detected phenomenon, thus machine learning approach was chosen for this task. Bidirectional Long Short-term Memory (BiLSTM) networks ^25^ have shown to be very successful for signal classification and regression tasks, thus it is suitable for the purpose of detection of cell death in analysed cell signals. The proposed approach is inspired by the detection of correlation filters ^26^, where the aim is to regress the Gaussian curve on the desired position, where Gaussian represent uncertainty in position. LSTM network has been trained for the regression of Gaussian curves created on the time of cell death (with sigma 50), where the whole method is summarized in Fig. 2.

**Figure 2.**
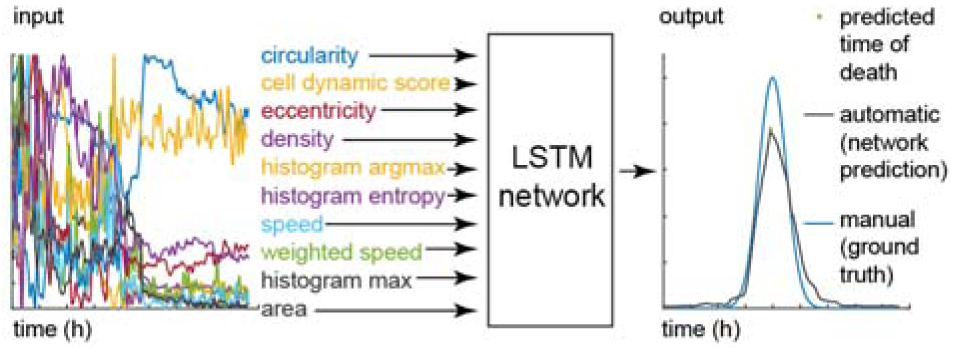
Prediction of cell death timepoint. The network input is the set of time-lapse signals. Ground truth Gaussian (used for training) is created from manual cell death timepoint labels. LSTM, Long Short-term Memory

According to ^27^, the network was set to two BiLSTM layers (with 100 units) and 3 fully connected layers (with 100, 50 and 100 neurons) with ReLu and dropout with probability 0.5. The whole network was optimized using Mean Square Error loss and ADAM optimizer ^28^ with learning rate 10^-3^, β_1_ = 0.9, β_2_ = 0.999, gradient clipping to norm 1 ^29^, weight decay 10^-3^, batch size 256 and 40 epochs. We augment our training dataset with random clipping (shortening each signal by at most 1/3 of length). One network was trained for all 9 cell lines/experiments evaluated by cross-validation, where one FOV from all experiments was used for testing and the rest for training. Cell death time was identified as a maximum in the network response with a value higher than the chosen threshold 0.4. Quality of cell detection is evaluated in terms of accuracy, where detection is considered as correct if it detects a death in a ±5h (±100 frames) window from ground truth cell death or if death is not detected and the cell is labelled as alive (at the end of the experiment).

### Cell death type identification

We observed two distinct dynamical patterns of cell mass manipulation during cellular death, where each cell was manually labelled as one of these types. To quantify these morphological types we extracted average values of extracted features in 200 frames (10 h) before cell death (shorter time window is used if more timepoints are not available). We trained linear Support Vector Machine (SVM) classifier for automatic classification of these cell death types. SVM classifier was trained for each cell line (one for all three treatments) because cell lines are morphologically different. All possible subsets of features were tested and classification accuracy was evaluated. Only density and CDS was finally used because accuracy does not increase significantly with the addition of other features (accuracy 75.4 % and 76.2 % for two and three features, respectively), where two features can be comprehensibly visualized. These two features have also a distinct biological meaning and its difference in an average aligned signal can be confirmed in Fig 3.

**Figure 3.**
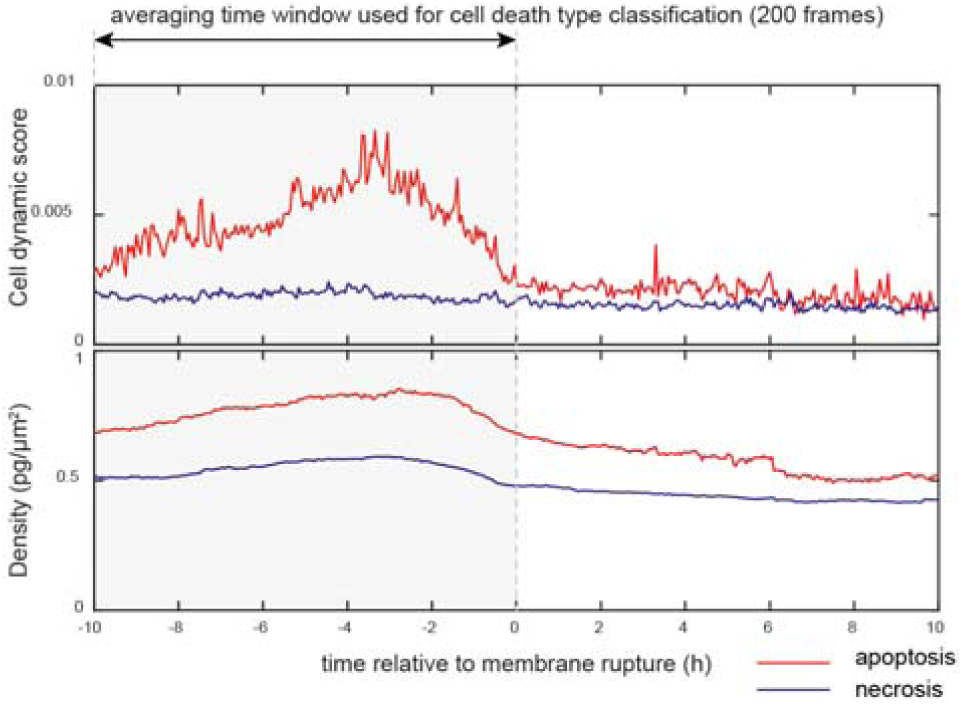
Classification of apoptosis and necrosis. The input parameters for this analysis are: (1) interval minus 10 h to timepoint of death (predicted in the previous step), and (2) manually annotated death type (apoptosis vs necrosis). Figure showing the average cell dynamic score and mass density, LNCaP cells, 0.1 μM doxorubicin treatment (N = 61 apoptosis and 53 necrosis).

**Figure 4.**
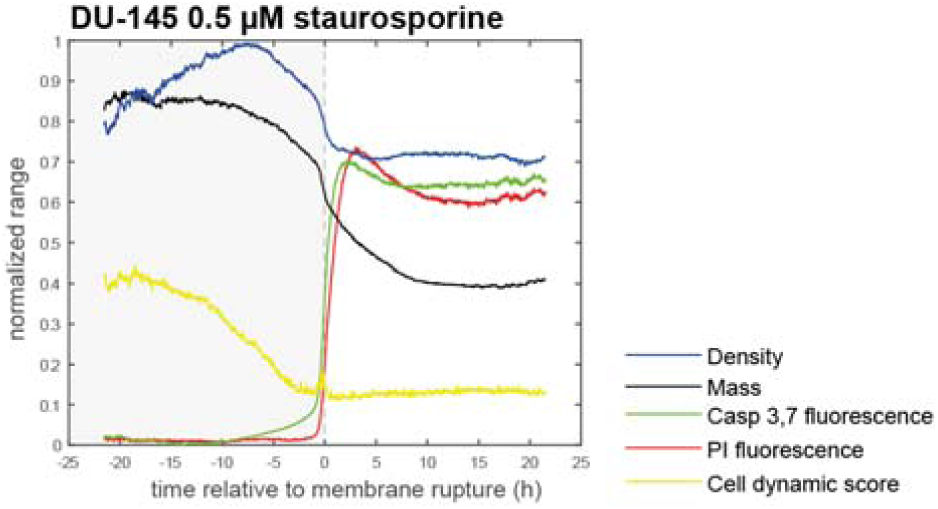
Necrotic cell death subroutine and its typical quantitative phase vs. fluorescence features. DU-145 cells, staurosporine treatment. Average curves of particular parameters for manually annotated N = 148 necrotic cells. PI, propidium iodide.

**Figure 5.**
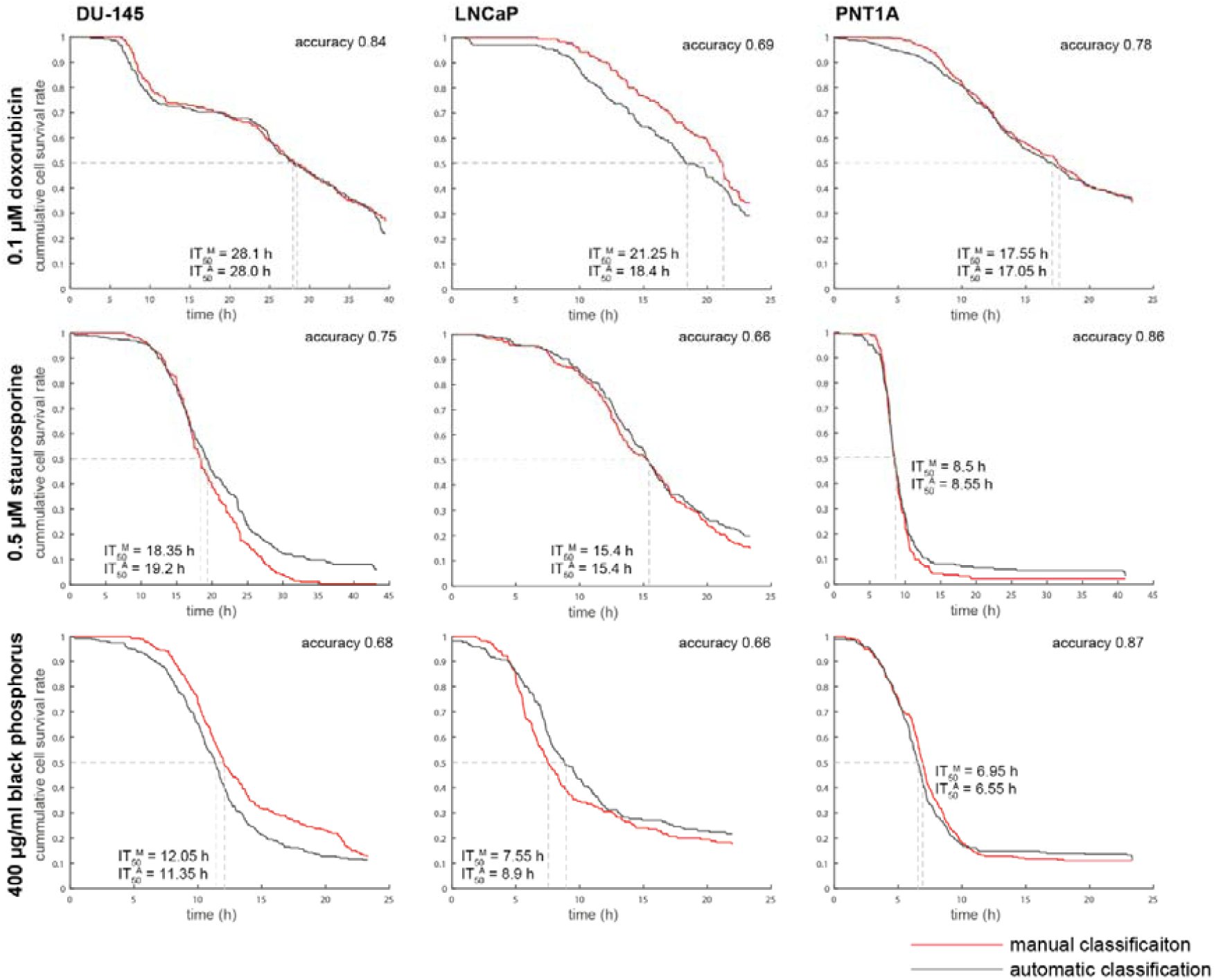
The cumulative survival rate of cells exposed to cell death inducers. Prediction of IT50 for cell populations. N(cells) = 651, 453, and 661 for DU-145, LNCaP, and PNT1A. Time-point of cell death for each cell was detected manually and automatically.

Furthermore, the correctness of the classification was confirmed by the analysis of nucleus shrinkage and by Casp 3,7–PI signal onset delay in fluorescence data. Cell shrinkage was measured as an average Hoechst 33342 brightness in 200 frames (10 h) before cell death (based on nuclei mask produced by GMM tracking). Casp 3,7–PI signal onset delay was measured as a time difference (delay) between the time when Casp 3,7 signals reach 1/3 of its average value after cell death (average in widnow of 200 frames) and the cell death time (which corresponds to a steep PI signal increase). The output of this analysis is shown in Fig.6C and D.

## Results

### Automatic cell death detection

After induction of cell death by using 0.5 µM staurosporine, 0.1 µM doxorubicin, or 400µg/mL of BP, respectively, 11 morphological parameters detectable by QPI (see Fig. 2) and their real-time values were collected. Effector caspase 3/7 activity, nuclear morphology and membrane integrity (propidium iodide signal; PI) were verified using time-lapse fluorescent microscopy to estimate ground truth for cell death detection based on cell mass parameters. The decrease in cell dry mass (pg), cell density (pg/pixel), and the Cell Dynamic Score (CDS; see the methodological part for equation) measured by QPI was in clear relationship with the onset of propidium iodide and caspase signalling. Furthermore, dead cells showed no longer any significant changes in CDS. (see Fig.4). Based on QPI features, automatic and label-free detection of IT_50_ (time of half-maximal inhibition effect for given treatment concentration) can be performed (see Fig.5.). Its evaluation is based on automatic cell detection of cell death using the LSTM network, where we can easily accumulate the number of dead cells assuming that all cells are alive at the start of the image sequence. This parameter is important because IC_50_ values can be significantly different in different time points (such as 24h versus 48h treatment ^30^). For IT_50_ of tested compounds and cell lines see Fig.5. Automatic detection of cell death showed 76% accuracy compared to the manual detection of cell death based on the fluorescent PI signal and morphological criteria visible to the naked eye.

### Cell density and Cell Dynamic Score reflect subroutine of cell death

Based on QPI data, we were able to distinguish two different subroutines of cell death: a) cells having high cell density and intensive blebbing of the plasma membrane (high CDS) followed by plasma membrane rupture and b) cells having low cell density and low CDS before plasma membrane rupture (see Fig. 7 and Supplementary video 1-2). These variants occupied extreme positions on the CDS versus cell density plot (see Fig.6b and Supplementary fig. 1). Time-lapse images and QPI features of these cells are shown in Fig 6a and Supplementary fig. 3 and 8. Based on fluorescent data referring caspase 3, 7 activity, nuclear morphology and membrane integrity (PI signal); see Fig.7 and Supplementary fig. 2-12, the a) type of cell death with high CDS and high cell density is *bona fide* apoptosis because PI signal was delayed over caspase signals and shrinkage of the nucleus was apparent. On the other hand, the b) type of cell death is *bona fide* necrosis as PI and caspase signals were displayed simultaneously and no significant shrinkage of the nucleus was apparent. Cells near the dividing line showed a)-like and b)-like features to varying degrees. Casp 3,7–PI signal onset delay and nucleus shrinkage (described in Methods) were quantified for both automatic and manual labels of cell death types (see Fig.6c and Supplementary table 1), which confirmed the existence of these two cell death subroutines. Furthermore, apoptosis and necrosis were visually detected by the expert according to the fluorescent signals and eye-visible cell morphology. The automatic label-free distinction of cell death subroutine showed 75.4 % accuracy (average of all cells and treatments) compared manual distinction of cell death type based on fluorescent signals and eye-visible morphological criteria; (see Fig.6e.). Interestingly, no cells showed a gradual cell rounding and loss of surface contact during staurosporine treatment. Other treatments caused such phenomena relatively often. For illustration see Fig.8.

**Figure 6.**
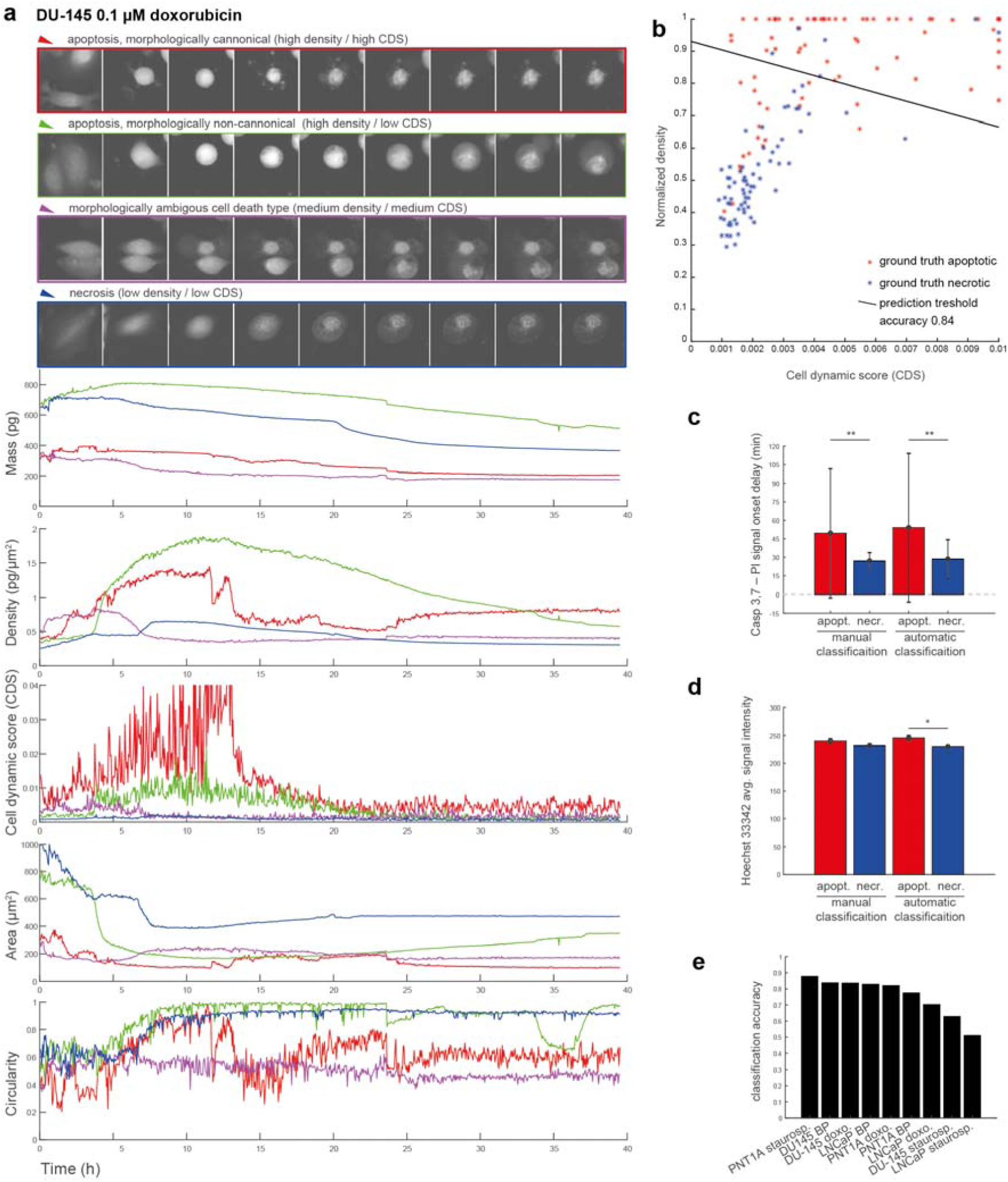
Quantitative phase-related parameters of cells dying by caspase 3, 7 dependent and independent cell death type. DU-145 cells exposed to 0.1 μM doxorubicin. **a.** Real-time QPI signals for canonical apoptosis (red), necrosis (blue), morphologically non-canonical apoptosis (green), and ambiguous cell death type (violet) based on a proposed classification algorithm. **b.** classification of cell death type according to cell dynamic score and normalized density. Cell death type was characterised by the delay of fluorescence onset of Casp 3,7 and PI signal **(c)**, and by different nuclear Hoechst 33342 signal intensity (**d)** * indicate p < 0.05, ** indicate p < 0.001**. e.** prediction accuracy for all other cell types exposed to other treatments (for results see Supplementary fig. 2 to 12). Bar charts are shown as mean and standard error. See Fig. 7. for quantitative phase and fluorescent images in critical timepoints of particular cells. BP, black phosphorus; PI, propidium iodide; CDS, cell dynamic score

**Figure 7.**
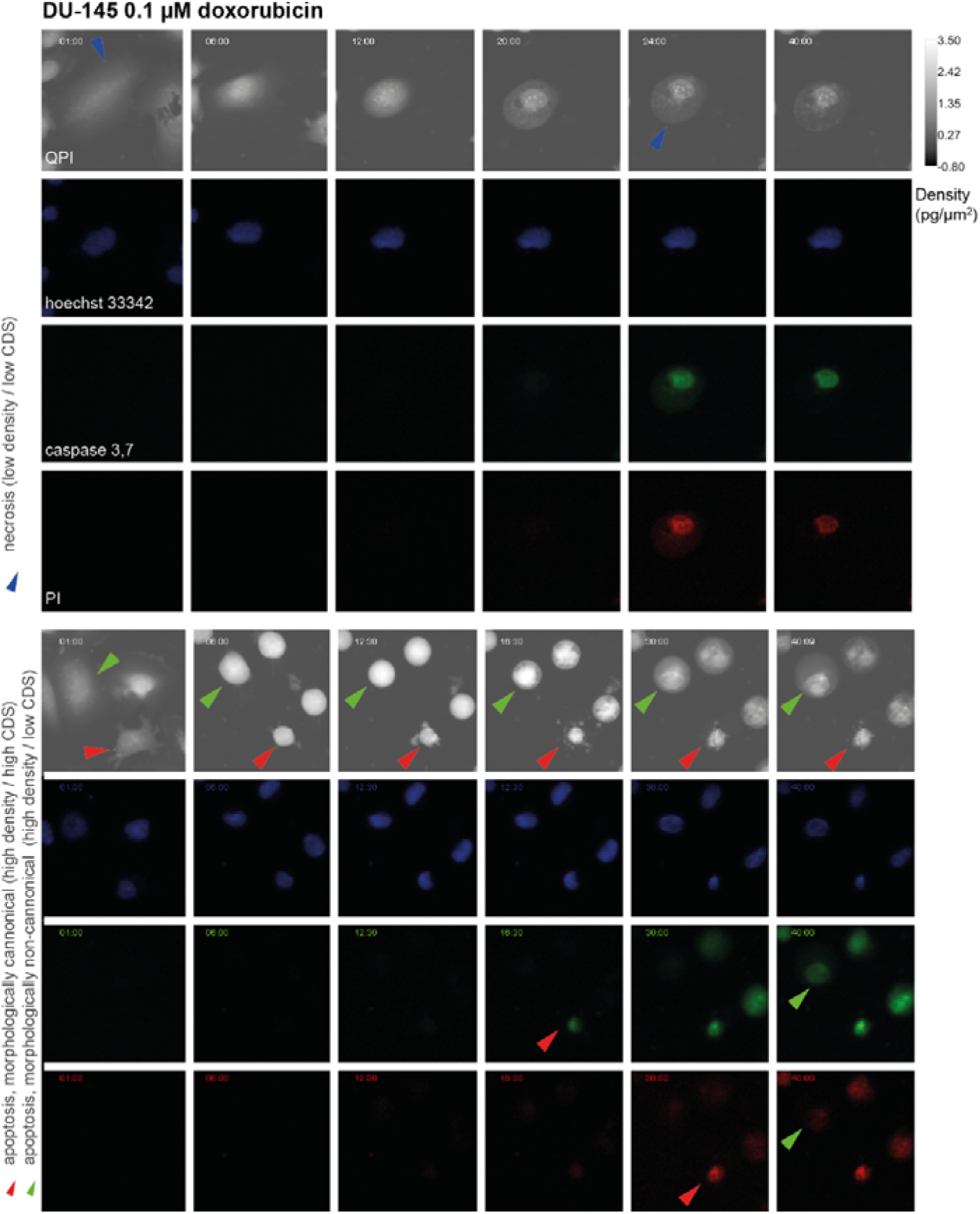
Characteristic quantitative phase data of necrotic and apoptotic cell (focused on typical morphological events). data correspond to time signals shown in fig. 6. Blue arrows indicate particular necrotic cell and membrane rupture, red (canonical apoptosis) and green (noncanonical apoptosis) arrows indicate particular apoptotic cells and their fluorescence onset of Casp 3,7 and PI signal. 10x magnification. FOV size approx. 107 μm. CDS, cell dynamic score; QPI, quantitative phase image; PI, propidium iodide.

**Figure 8.**
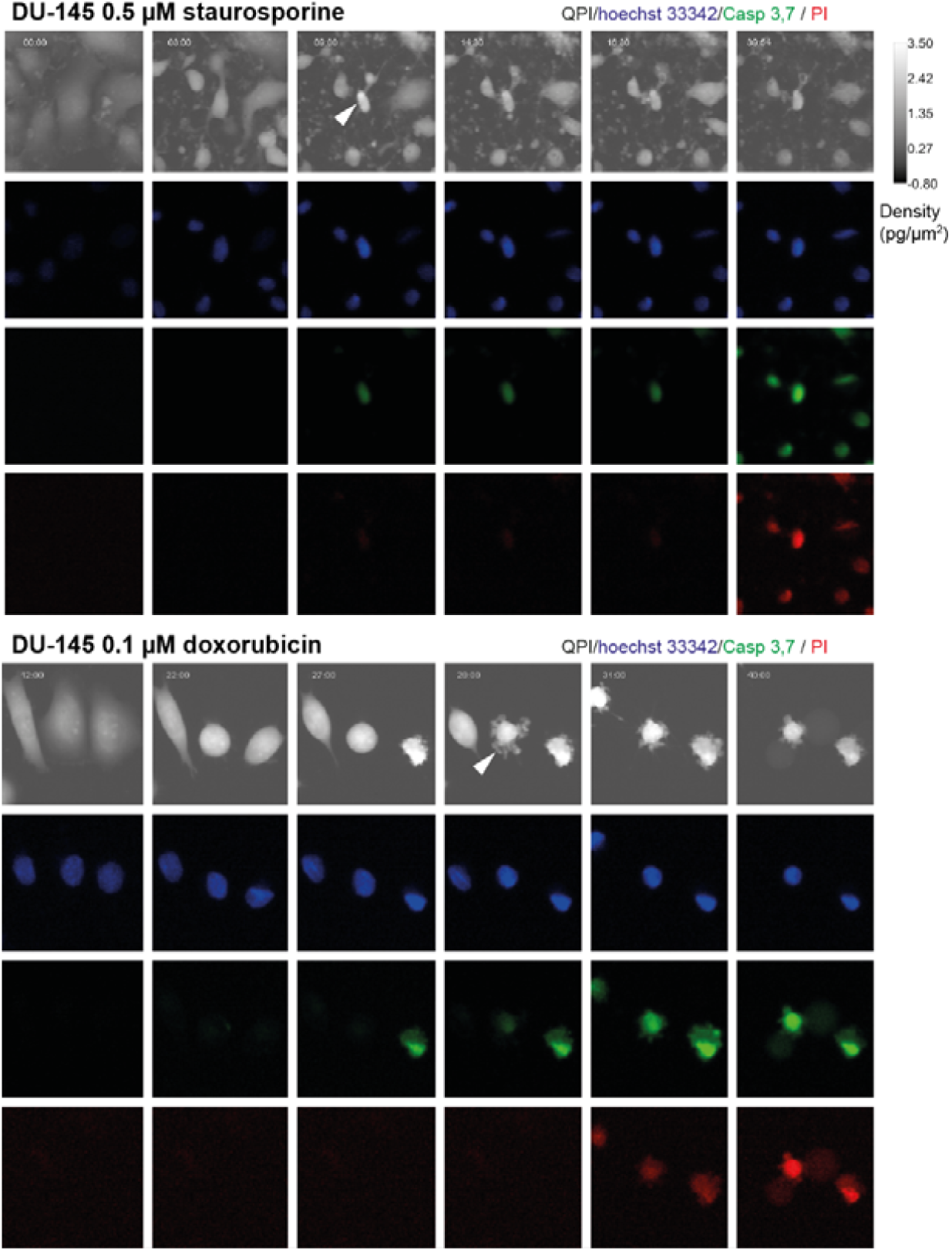
Quantitative phase image of staurosporine- and doxorubicin-induced apoptosis; morphological differences. White arrows indicate persisting attachment of staurosporine-treated cells and surface detachment of doxorubicin-treated ones. 10x magnification. FOV size approx. 107 μm. QPI, quantitative phase image; PI, propidium iodide.

## Discussion

The basic requirement for a good method of cell death detection is the ability to detect a robust parameter in a highly reproducible and if possible inexpensive manner under changing conditions caused by the cell heterogeneity. In this study, we suggest that morphological and dynamical features extracted from the QPI cell image are suitable for cell death detection. QPI enables the time-lapse observation of subtle changes in the cell mass distribution unrecognizable by the naked eye, and therefore provides detailed information on cell morphology and cell mass topography during cell death. Cell dry mass can be calculated directly from phase values detected in each pixel ^31, 32, 33^. Cell dry mass is the direct result of biosynthetic and degradative processes within a cell and is, therefore, a promising indicator of cell fate including cell death ^34, 35^. The distribution of cell mass during the processes of cell death is significantly changing in time with a typical steep decrease of the cell mass due to the rupture of the plasma membrane or cell fragmentation. This phenomenon *is bona fide* universal for all cell types and was also observed in target cells after contact with a cytotoxic T-cells ^34^. Plasma membrane rupture can be otherwise measured by quantification of the release of intracellular enzymes from the cell into the cell culture medium. Nevertheless, the enzymatic activity of these enzymes can be seriously affected by differences in the pH between intracellular environment and culture medium and by time spent in the extracellular space ^3^.

Based on QPI data, we were also able to distinguish two specific subroutines of cell death. While the cell death subroutine a) (*bona fide* apoptosis) showed a high cell density and strong fluctuation in CDS before cell death; during the subroutine b) (*bona fide* necrosis) the cell density was low and CDS was relatively stable after the initial decline. These variants occupied extreme positions on the CDS versus cell density plot. Cells near the dividing line (see Fig.6b.) probably succumb to another subroutine of cell death such as pyroptosis or necroptosis, as neither staurosporine nor doxorubicin is a specific apoptosis inductor ^36, 37^. The a) type of cell death was characteristic by a decrease in the cell area, a gradual cell rounding and a loss of surface contact (this phenomenon was not observed in the case of staurosporine treatment) followed by membrane blebbing (known as “dance of death”); see Fig. 6 and Supplementary fig. 2 to 12, ^38^. Since membrane blebbing during apoptosis results from caspase-mediated activation of ROCK I, it can be assumed that apoptosis was involved in these cases ^39^. At the final stages of apoptosis, the actin cytoskeleton is degraded ^40, 41^. Due to the hydrophobic nature of the apoptotic bodies, they undergo the plasma membrane fusion in the culture medium. The membrane of this post-apoptotic body subsequently cracks and culminates in secondary necrosis (in our case depicted by a steep decrease of cell mass) in the absence of phagocytes ^42, 43^. The b) type of observed cell death presented persisting large cytoplasmic membrane blebs or multiple small blebs and cell swelling leading to the final cell membrane rupture. This type of dying cells was adherent during the whole process. It can be assumed that necrosis was involved in these cases ^3^. Although several studies based on QPI detection of cell death have been published earlier, they do not include the distinguishing of specific subroutines of cell death and have failed to capture the entire process of cell death, including the early stages due to short intervals of QPI capturing ^44, 45, 46^. From an image analysis point of view, we introduce a new powerful technique for cell tracking and segmentation, capable of robustly track cell thought the whole image sequence. We also introduced a completely new method for the detection of cell death in time-series of measured cell features using LSTM neural network. To the best of our knowledge, this is also the first study introducing CDS and cell density as a parameter typical for individual cell death subroutines.

## Supporting information

Description of additional supplemetary files

## Data availability

The Matlab code is available at GitHub (https://github.com/tomasvicar/CellDeathDetect). Annotated quantitative phase image dataset (with cell tracking masks and labels of cell death timepoints and types of death) used in the manuscript is available at Zenodo repository (www.zenodo.org), https://doi.org/10.5281/zenodo.2601562

## Authors contributions

Tomas Vicar: image analysis, data interpretation, manuscript preparation; Martina Raudenska: wrote the manuscript, study design; Jaromir Gumulec: interpretation of results, manuscript preparation; Michal Masarik: study design, study concepts; Jan Balvan: study design, microscope operation, interpretation of results, writing the manuscript

## Acknowledgement

This work was supported by funds from the Faculty of Medicine, Masaryk University to Junior researcher (Jan Balvan), and by Grant Agency of the Czech Republic (18–24089 S). Black phosphorus was kindly provided by Dr. Martin Pumera and Dr. Zdenek Sofer from University of Chemistry and Technology Prague, Czech republic. We acknowledge Dr. Michaela Fojtu for her kind assistance with black phosphorus preparation and Tescan, a.s. for support to Tescan Q-PHASE technology.

## References

1. Galluzzi L, et al. Molecular mechanisms of cell death: recommendations of the Nomenclature Committee on Cell Death 2018. Cell Death Differ 25, 486–541 (2018).

2. Kroemer G, et al. Classification of cell death: recommendations of the Nomenclature Committee on Cell Death 2009. Cell Death and Differentiation 16, 3–11 (2009).

3. Galluzzi L, et al. Guidelines for the use and interpretation of assays for monitoring cell death in higher eukaryotes. Cell Death and Differentiation 16, 1093–1107 (2009).

4. Kepp O, Galluzzi L, Lipinski M, Yuan JY, Kroemer G. Cell death assays for drug discovery. Nature Reviews Drug Discovery 10, 221–237 (2011).

5. Balvan J, et al. Multimodal holographic microscopy: distinction between apoptosis and oncosis. PLoS One 10, e0121674 (2015).

6. Palchaudhuri R, et al. A Small Molecule that Induces Intrinsic Pathway Apoptosis with Unparalleled Speed. Cell reports 13, 2027–2036 (2015).

7. Yang F, Teves SS, Kemp CJ, Henikoff S. Doxorubicin, DNA torsion, and chromatin dynamics. Biochimica Et Biophysica Acta-Reviews on Cancer 1845, 84–89 (2014).

8. Fojtů M, et al. Black Phosphorus Cytotoxicity Assessments Pitfalls: Advantages and Disadvantages of Metabolic and Morphological Assays. Chemistry – A European Journal 25, 349–360 (2019).

9. Ulman V, et al. An objective comparison of cell-tracking algorithms. Nature Methods 14, 1141 (2017).

10. Tinevez J-Y, et al. TrackMate: An open and extensible platform for single-particle tracking. Methods 115, 80–90 (2017).

11. Huh S, Kanade T. Apoptosis detection for non-adherent cells in time-lapse phase contrast microscopy. Med Image Comput Comput Assist Interv 16, 59–66 (2013).

12. Manivannan S, Li W, Akbar S, Wang R, Zhang J, McKenna SJ. HEp-2 Cell Classification Using Multi-resolution Local Patterns and Ensemble SVMs. In: 2014 1st Workshop on Pattern Recognition Techniques for Indirect Immunofluorescence Images (ed^(eds) (2014).

13. Bs D, Subramaniam K, H R N. HEp-2 cell classification using artificial neural network approach (2016).

14. Li H. Deep CNNs for HEp-2 Cells Classification : A Cross-specimen Analysis (2016).

15. Skjoth IH, Issinger OG. Profiling of signaling molecules in four different human prostate carcinoma cell lines before and after induction of apoptosis. Int J Oncol 28, 217–229 (2006).

16. Mitchell S, Abel P, Ware M, Stamp G, Lalani E. Phenotypic and genotypic characterization of commonly used human prostatic cell lines. BJU Int 85, 932–944 (2000).

17. Raudenska M, et al. Cisplatin enhances cell stiffness and decreases invasiveness rate in prostate cancer cells by actin accumulation. Scientific Reports 9, 1660 (2019).

18. Chappell WH, Lehmann BD, Terrian DM, Abrams SL, Steelman LS, McCubrey JA. p53 expression controls prostate cancer sensitivity to chemotherapy and the MDM2 inhibitor Nutlin-3. Cell cycle (Georgetown, Tex) 11, 4579–4588 (2012).

19. Prescher JA, Bertozzi CR. Chemistry in living systems. Nature Chemical Biology 1, 13–21 (2005).

20. Park Y, Depeursinge C, Popescu G. Quantitative phase imaging in biomedicine. Nature Photonics 12, 578–589 (2018).

21. Jung C, Kim C, Chae SW, Oh S. Unsupervised Segmentation of Overlapped Nuclei Using Bayesian Classification. IEEE Transactions on Biomedical Engineering 57, 2825–2832 (2010).

22. Nistér D, Stewénius H. Linear Time Maximally Stable Extremal Regions (2008).

23. Vicar T. Robust Cell Nuclei Tracking Using Gaussian Mixture Shape Model. In: 24th Conference STUDENT EEICT 2018 (ed^(eds). Brno University of Technology, Faculty of Electrical Engineering and Communication (2018).

24. Pinidiyaarachchi A, Wählby C. Seeded watersheds for combined segmentation and tracking of cells. In: International Conference on Image Analysis and Processing (ed^(eds). Springer (2005).

25. Graves A, Mohamed A, Hinton G. Speech recognition with deep recurrent neural networks. In: 2013 IEEE International Conference on Acoustics, Speech and Signal Processing (ed^(eds) (2013).

26. Henriques JF, Caseiro R, Martins P, Batista J. High-Speed Tracking with Kernelized Correlation Filters. IEEE Trans Pattern Anal Mach Intell 37, 583–596 (2015).

27. Reimers N, Gurevych I. Optimal hyperparameters for deep lstm-networks for sequence labeling tasks. arXiv preprint arXiv:170706799, (2017).

28. Kingma DP, Ba J. Adam: A method for stochastic optimization. arXiv preprint arXiv:14126980, (2014).

29. Pascanu R, Mikolov T, Bengio Y. On the difficulty of training recurrent neural networks. In: International conference on machine learning (ed^(eds) (2013).

30. Gumulec J, et al. Cisplatin-resistant prostate cancer model: Differences in antioxidant system, apoptosis and cell cycle. International Journal of Oncology 44, 923–933 (2014).

31. Slabý T, et al. Coherence-controlled holographic microscopy for live-cell quantitative phase imaging. (ed^(eds) (2015).

32. Chmelik R, et al. Chapter 5 - The Role of Coherence in Image Formation in Holographic Microscopy. In: Progress in Optics (ed^(eds Emil W). Elsevier (2014).

33. Kolman P, Chmelik R. Coherence-controlled holographic microscope. Optics Express 18, 21990–22003 (2010).

34. Zangle TA, Burnes D, Mathis C, Witte ON, Teitell MA. Quantifying biomass changes of single CD8+ T cells during antigen specific cytotoxicity. PLoS One 8, e68916 (2013).

35. Popescu G, et al. Optical imaging of cell mass and growth dynamics. American Journal of Physiology-Cell Physiology 295, C538–C544 (2008).

36. Simenc J, Lipnik-Stangelj M. Staurosporine induces different cell death forms in cultured rat astrocytes. Radiology and oncology 46, 312–320 (2012).

37. Sugimoto K, Tamayose K, Sasaki M, Hayashi K, Oshimi K. Low-dose doxorubicin-induced necrosis in Jurkat cells and its acceleration and conversion to apoptosis by antioxidants. Br J Haematol 118, 229–238 (2002).

38. Galluzzi L, et al. Molecular definitions of cell death subroutines: recommendations of the Nomenclature Committee on Cell Death 2012. Cell Death and Differentiation 19, 107–120 (2012).

39. Coleman ML, Sahai EA, Yeo M, Bosch M, Dewar A, Olson MF. Membrane blebbing during apoptosis results from caspase-mediated activation of ROCK I. Nat Cell Biol 3, 339–345 (2001).

40. Coleman ML, Olson MF. Rho GTPase signalling pathways in the morphological changes associated with apoptosis. Cell Death Differ 9, 493–504 (2002).

41. Desouza M, Gunning PW, Stehn JR. The actin cytoskeleton as a sensor and mediator of apoptosis. Bioarchitecture 2, 75–87 (2012).

42. Silva MT. Secondary necrosis: the natural outcome of the complete apoptotic program. FEBS Lett 584, 4491–4499 (2010).

43. Hanahan D, Weinberg RA. Hallmarks of Cancer: The Next Generation. Cell 144, 646–674 (2011).

44. Pavillon N, et al. Early Cell Death Detection with Digital Holographic Microscopy. PLoS One 7, (2012).

45. Khmaladze A, Matz RL, Epstein T, Jasensky J, Holl MMB, Chen Z. Cell volume changes during apoptosis monitored in real time using digital holographic microscopy. Journal of Structural Biology 178, 270–278 (2012).

46. Pavillon N, et al. Cell morphology and intracellular ionic homeostasis explored with a multimodal approach combining epifluorescence and digital holographic microscopy. Journal of Biophotonics 3, 432–436 (2010).

