## Supplementary material for "Detection and characterization of apoptotic and necrotic cell death by time-lapse quantitative phase image analysis": Description of additional supplemetary files

Description of additional supplementary files

**Supplementary video 1:** **Characteristic quantitative phase data and corresponding signals of necrotic cell (low cell density, low cell dynamic score).** DU-145 cells exposed to 0.1 μM doxorubicin. Vertical red line indicate membrane rupture. 10x magnification. CDS, cell dynamic score; QPI, quantitative phase image; PI, propidium iodide.

**Supplementary video 2:** **Characteristic quantitative phase data and corresponding signals of morphologically canonical apoptosis (high cell density, high cell dynamic score).** DU-145 cells exposed to 0.1 μM doxorubicin. Vertical red line indicate membrane rupture. 10x magnification. CDS, cell dynamic score; QPI, quantitative phase image; PI, propidium iodide.
