## Supplementary material for "Detection and characterization of apoptotic and necrotic cell death by time-lapse quantitative phase image analysis": Supplemetary information

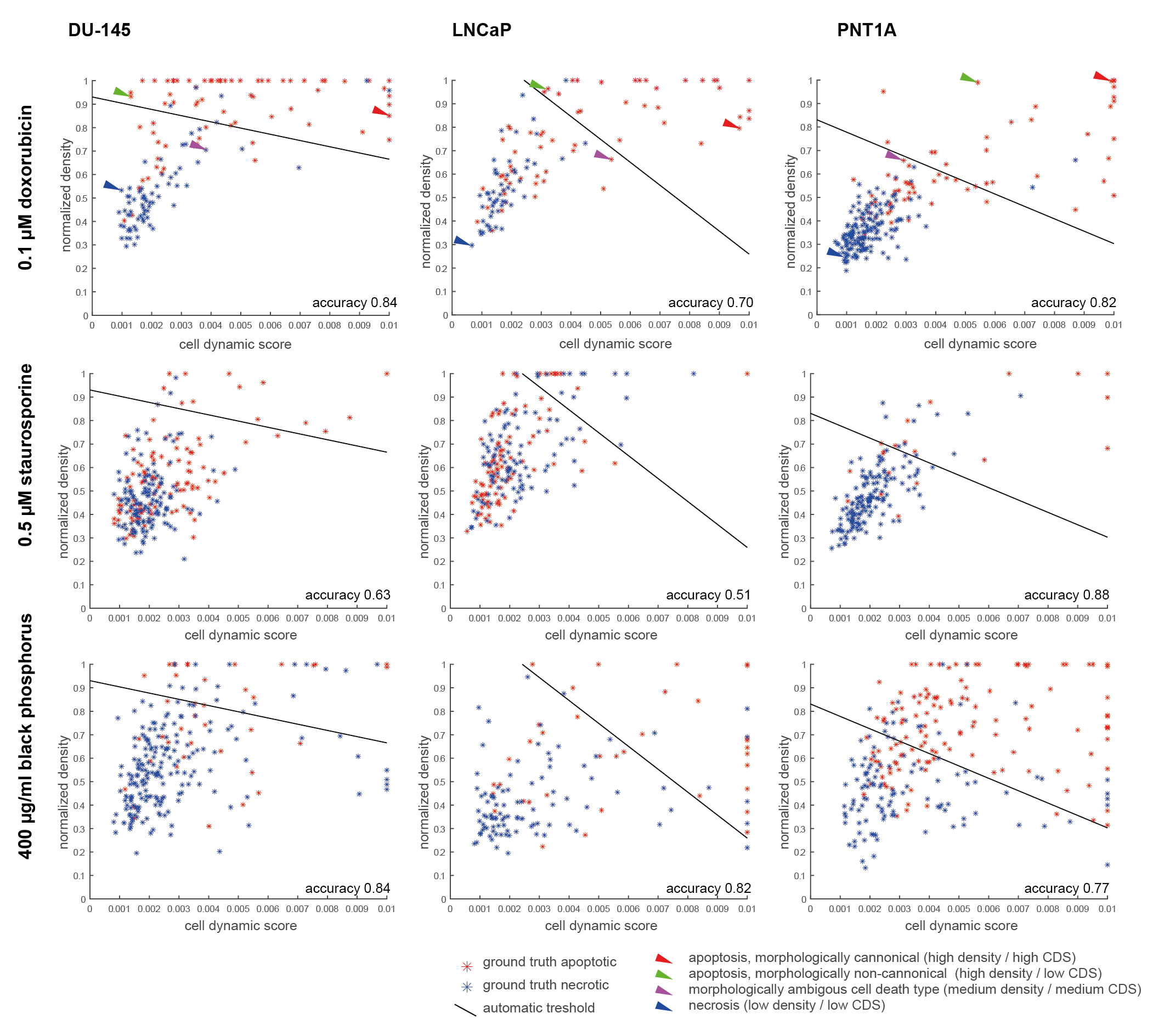

**Supplementary figure 1: Classification of cell death type according to cell dynamic score and normalized density.** Line indicates automatic prediction threshold (based on linear SVM), red and blue points indicate manually annotated apoptosis and necrosis. Arrows indicate morphologically typical examples shown in Supplementary fig. 2 to 12. CDS, cell dynamic score.

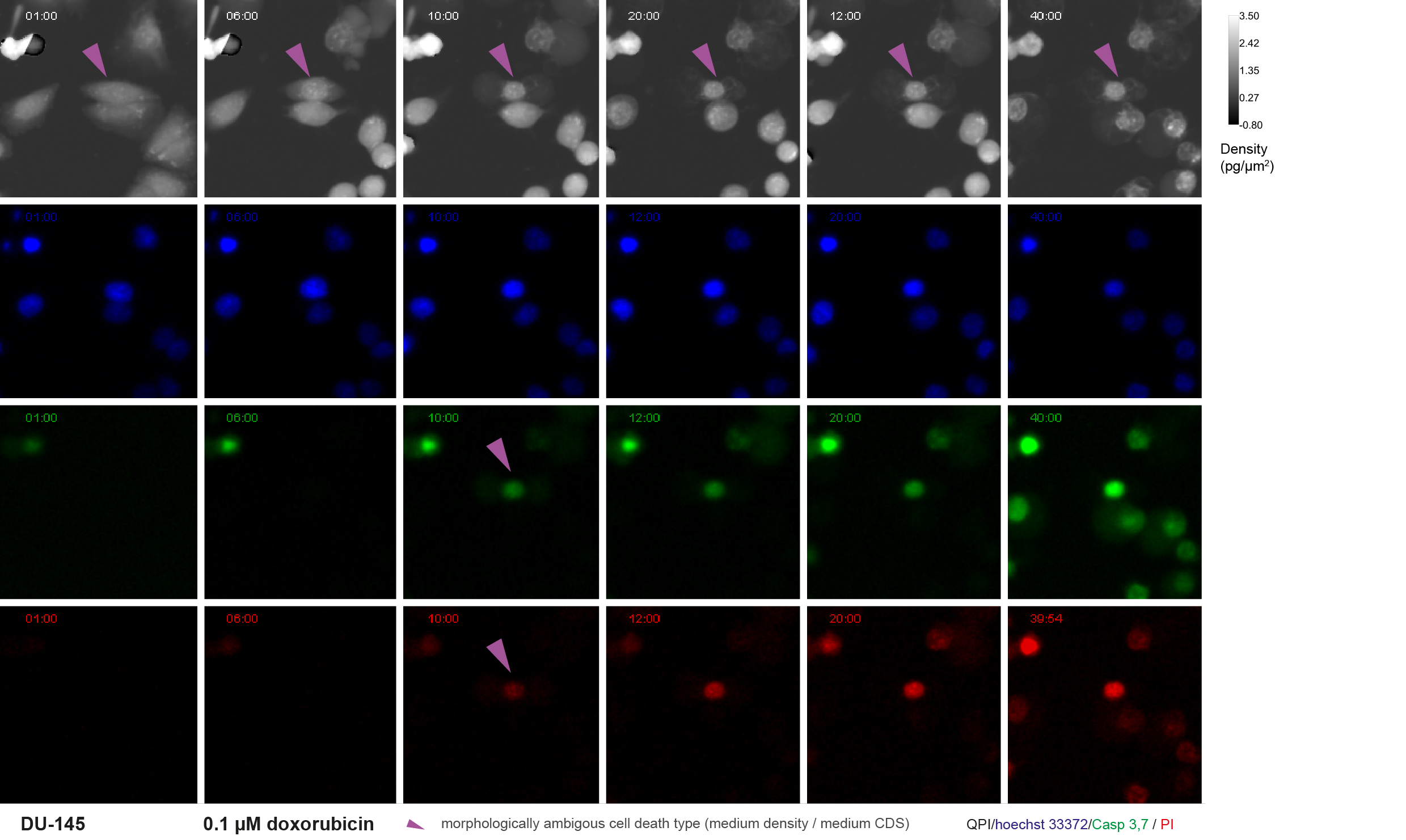

**Supplementary fig. 2: Critical timepoints for ambiguous cell death - quantitative phase and fluorescence image (medium cell density – medium cell dynamic score)**. DU-145 cells exposed to 0.1 μM doxorubicin. Violet arrows indicate analysed cell undergoing ambiguous death type (QPI image) and membrane rupture (fluorescence image). For quantitative time-lapse phase-related signals see Fig. 6. 10x magnification. FOV size approx. 107 μm. QPI, quantitative phase image; PI, propidium iodide.

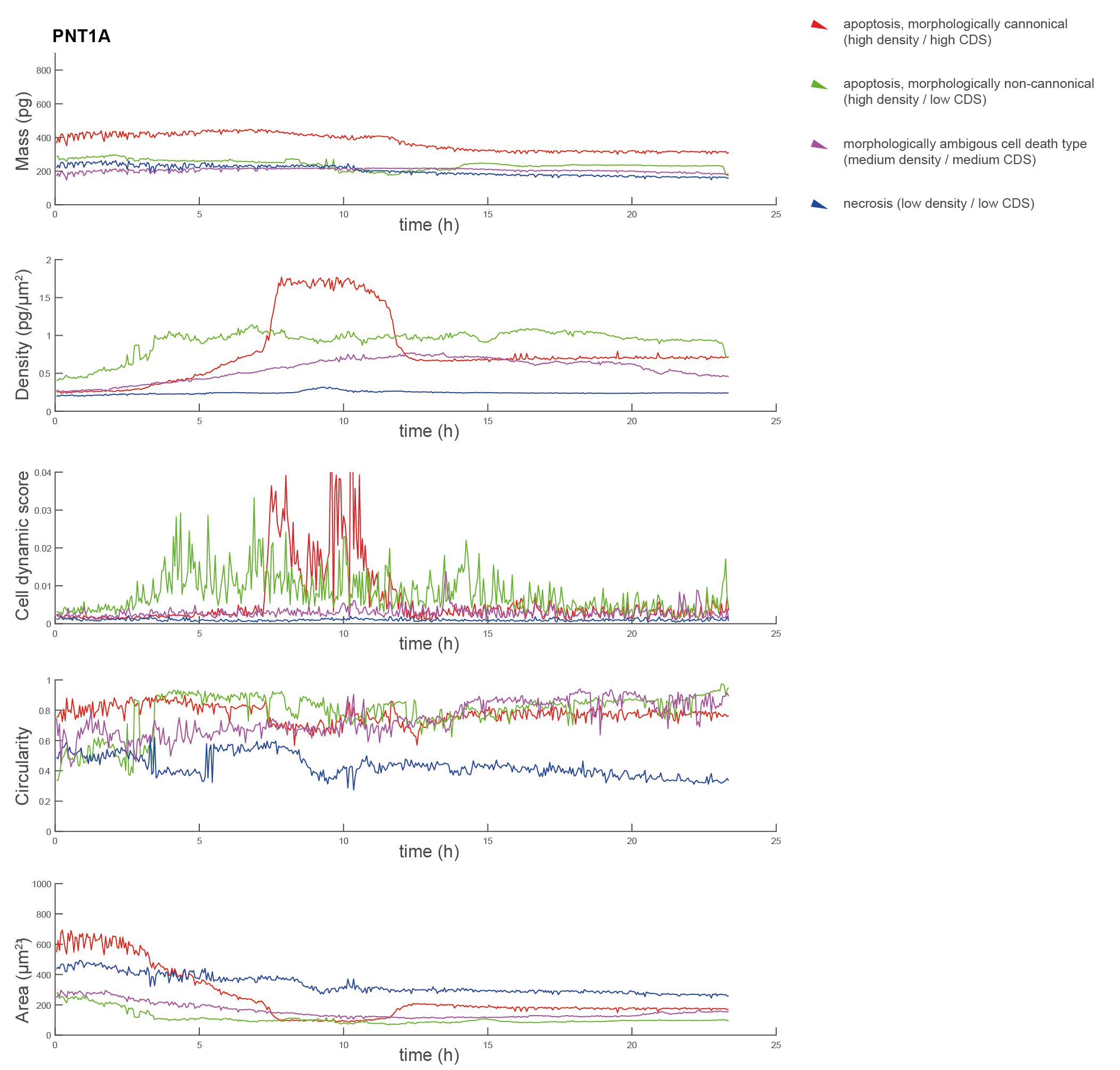

**Supplementary fig. 3: PNT1A cells Quantitative phase-related parameters of morphologically representative cell death type.** Cells exposed to 0.1 μM doxorubicin. Time signals for QPI parameters for morphologically canonical apoptosis (red), necrosis (blue), morphologically non-canonical apoptosis (green), and ambiguous cell death type (violet) based on a proposed classification algorithm. For examples of characteristic cells see Supplementary fig 4 to 7. CDS, cell dynamic score

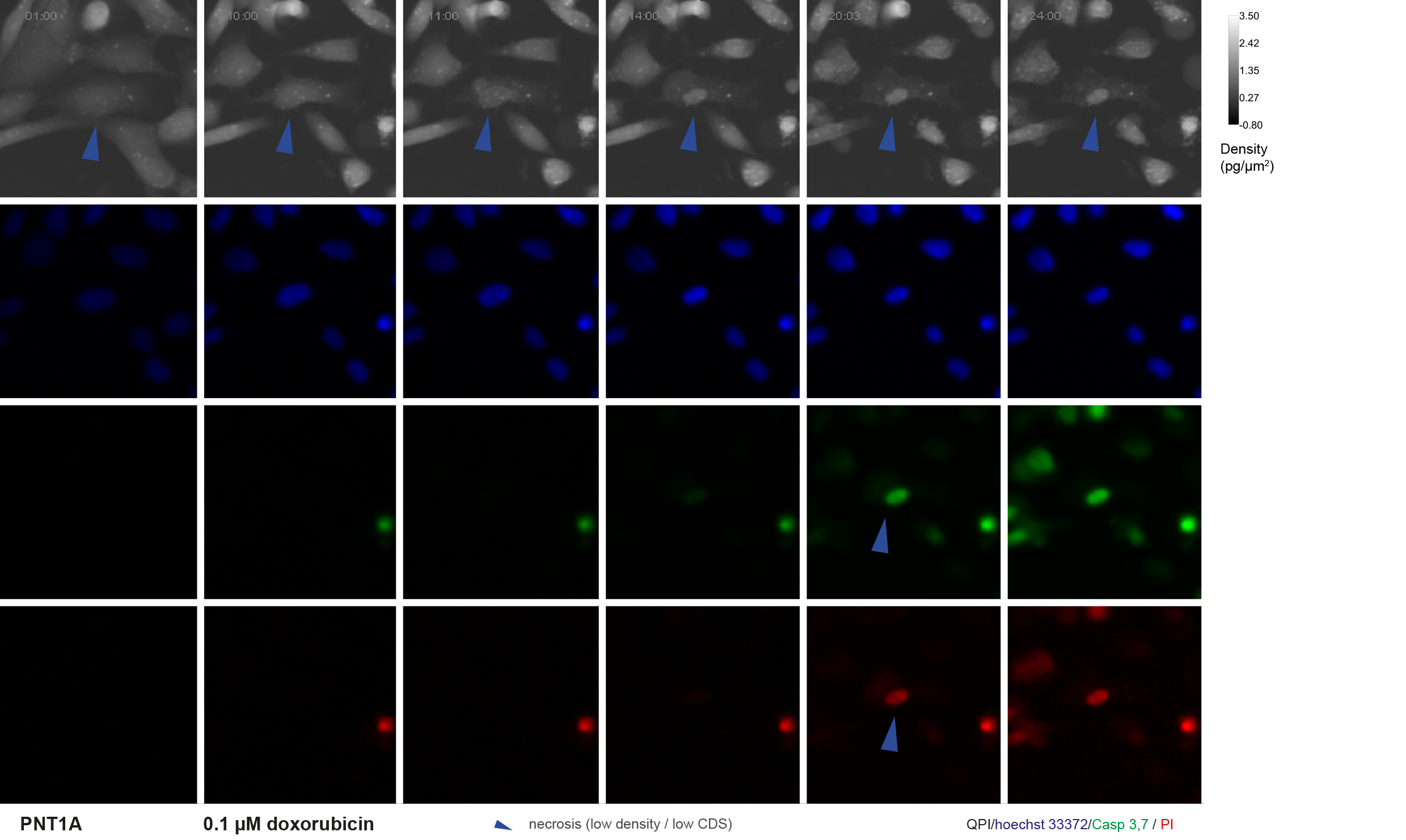

**Supplementary fig. 4:** **Critical timepoints for necrotic cell death - quantitative phase and fluorescence image (low cell density – low cell dynamic score)**. PNT1A cells exposed to 0.1 μM doxorubicin. Blue arrows indicate analysed necrotic cell (QPI image) and membrane rupture (fluorescence image). Notice simultaneous onset of PI and Casp 3, 7 signals. For quantitative time-lapse phase-related signals see supplementary fig. 3. 10x magnification. FOV size approx. 107 μm. QPI, quantitative phase image; PI, propidium iodide.

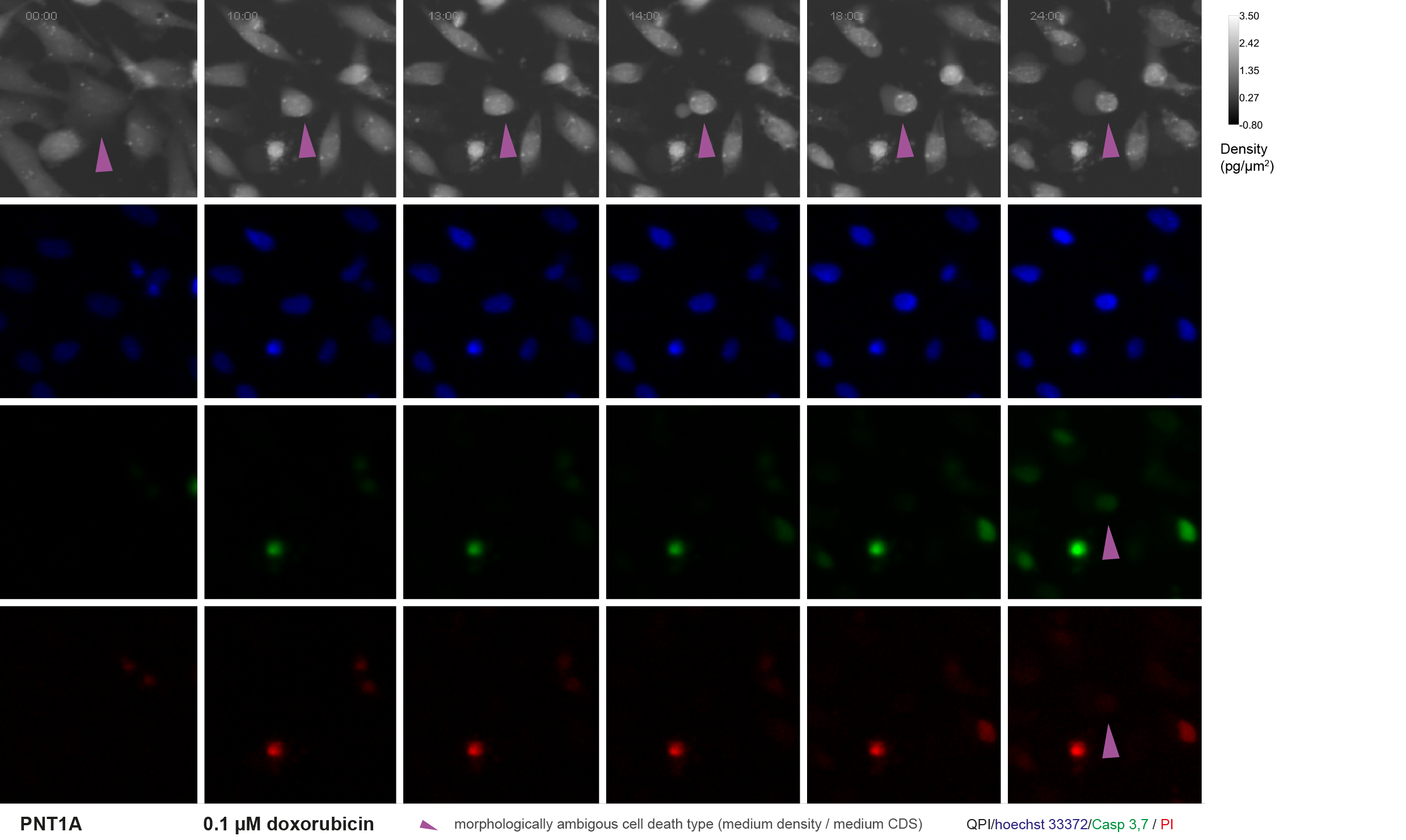

**Supplementary fig. 5: Critical timepoints for ambiguous cell death - quantitative phase and fluorescence image (medium cell density – medium cell dynamic score)**. PNT1A cells exposed to 0.1 μM doxorubicin. Violet arrows indicate analysed cell undergoing ambiguous death type (QPI image) and membrane rupture (fluorescence image). For quantitative time-lapse phase-related signals see supplementary fig. 3. 10x magnification. FOV size approx. 107 μm. QPI, quantitative phase image; PI, propidium iodide.

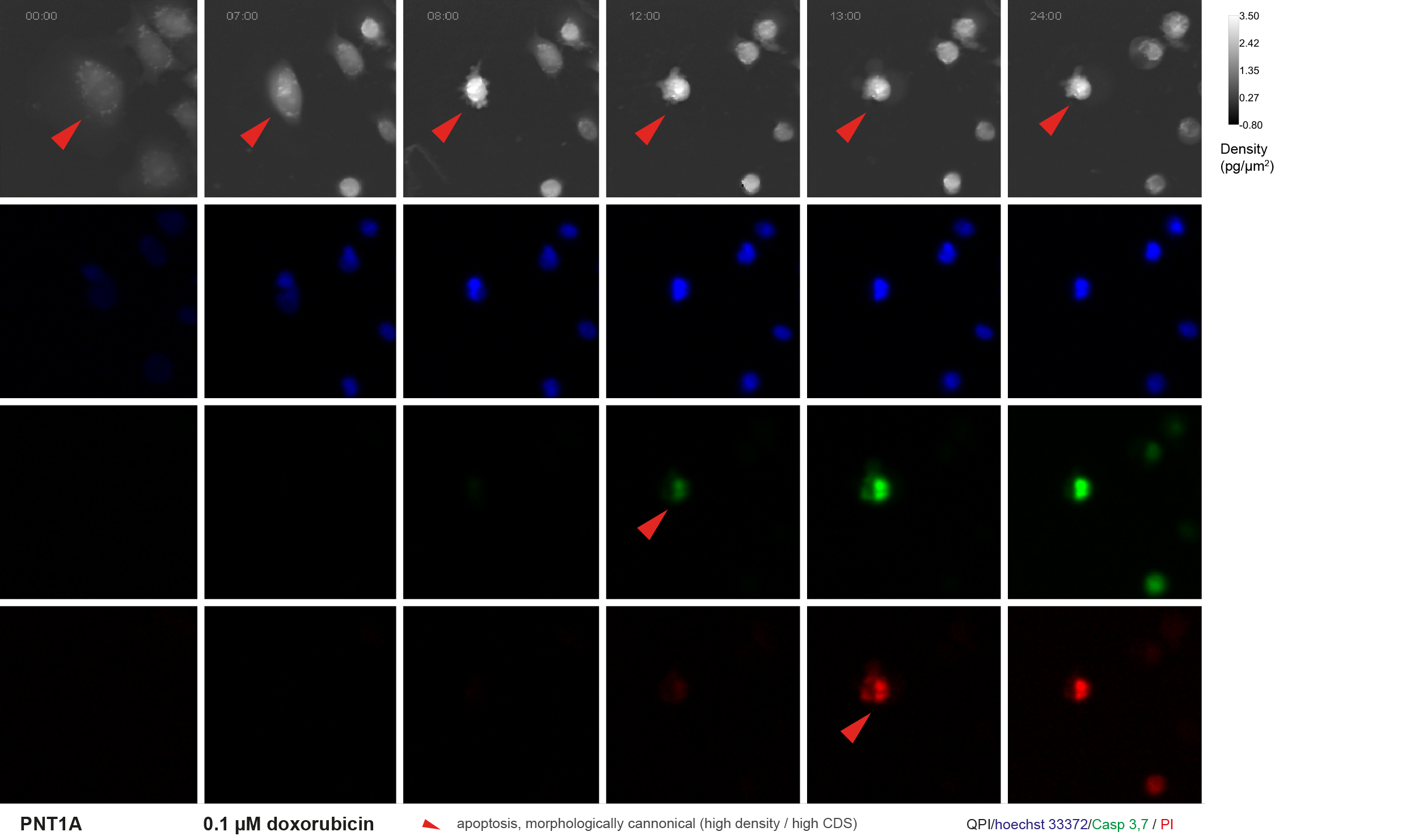
**Supplementary fig. 6: Critical timepoints for morphologically canonical apoptosis - quantitative phase and fluorescence image** **(high cell density – high cell dynamic score)**. PNT1A cells exposed to 0.1 μM doxorubicin. Red arrows indicate analysed cell undergoing cannonical apoptosis (QPI image) and membrane rupture (fluorescence image). Notice the onset of Casp 3,7 fluorescence before PI. For quantitative time-lapse phase-related signals see supplementary fig. 3. 10x magnification. FOV size approx. 107 μm. QPI, quantitative phase image; PI, propidium iodide.

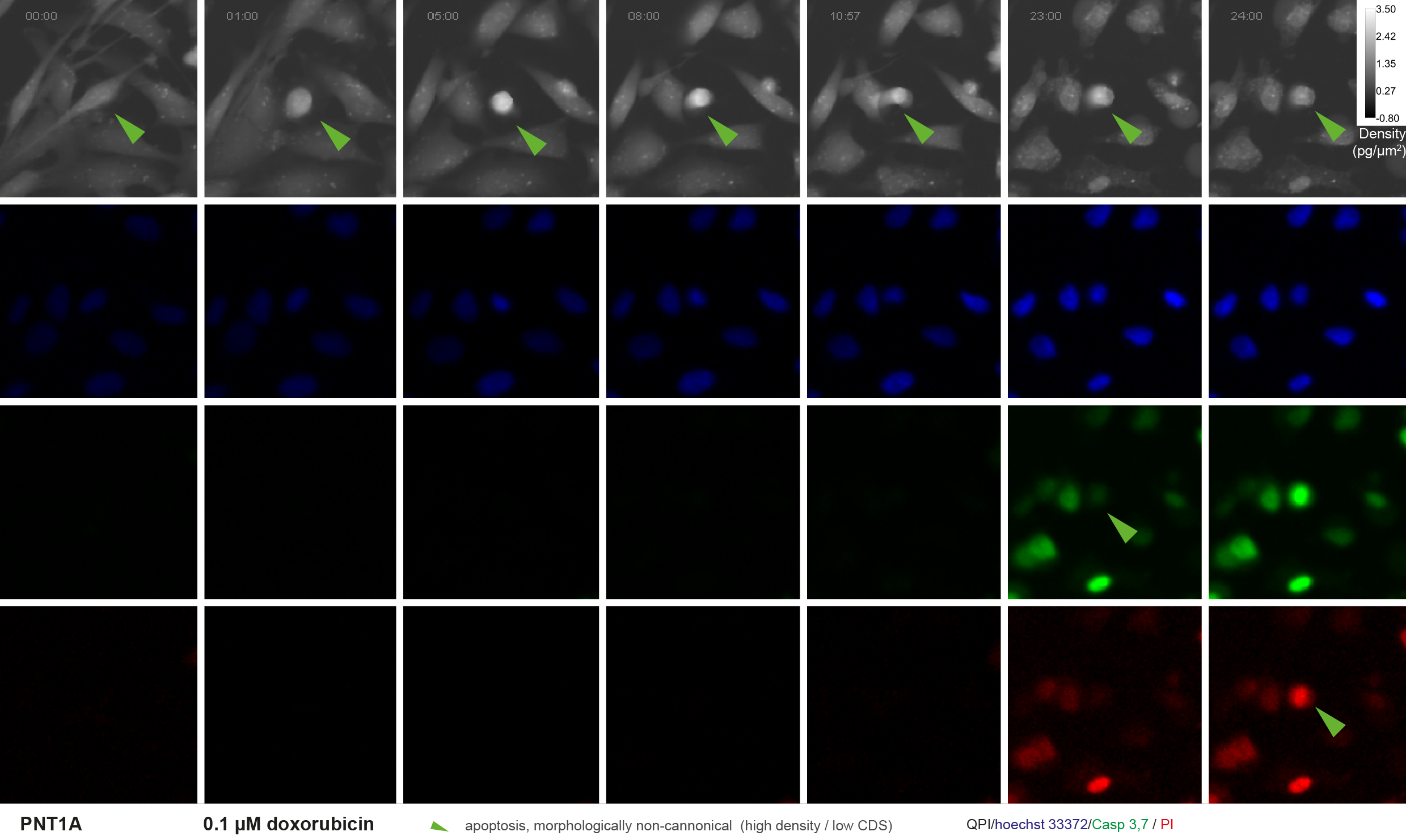

**Supplementary fig.7: Critical timepoints for morphologically non-canonical apoptosis - quantitative phase and fluorescence image**. PNT1A cells exposed to 0.1 μM doxorubicin. Red arrows indicate analysed cell undergoing cannonical apoptosis (QPI image) and membrane rupture (fluorescence image). For quantitative time-lapse phase-related signals see supplementary fig. 3. 10x magnification. FOV size approx. 107 μm. QPI, quantitative phase image; PI, propidium iodide.

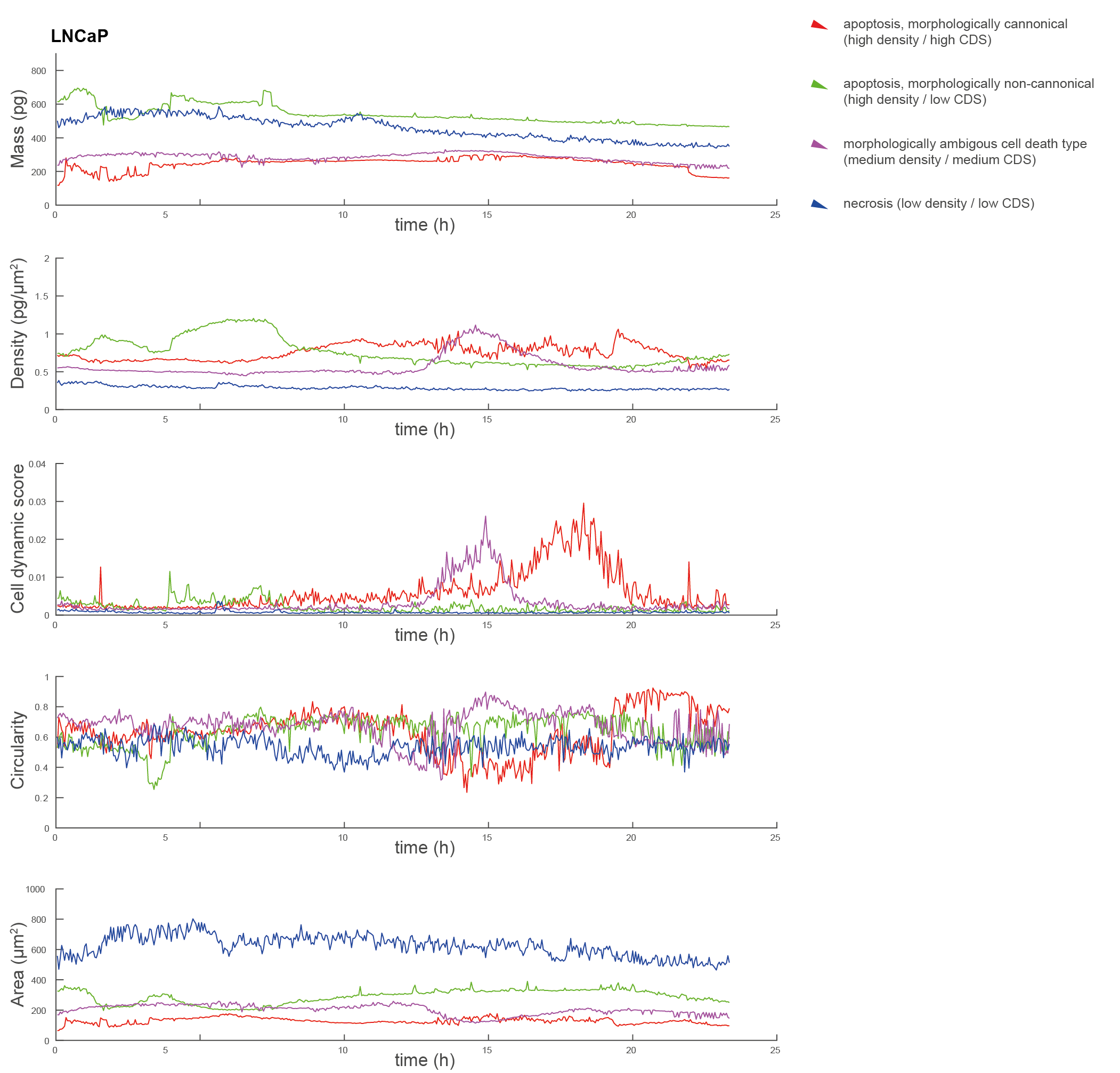

**Supplementary fig. 8: LNCaP cells Quantitative phase-related parameters of morphologically representative cell death type.** Cells exposed to 0.1 μM doxorubicin. Time signals for particular parameters for morphologically canonical apoptosis (red), necrosis (blue), morphologically non-canonical apoptosis (green), and ambiguous cell death type (violet) based on a proposed classification algorithm. CDS, cell dynamic score; QPI, quantitative phase image; PI, propidium iodide.

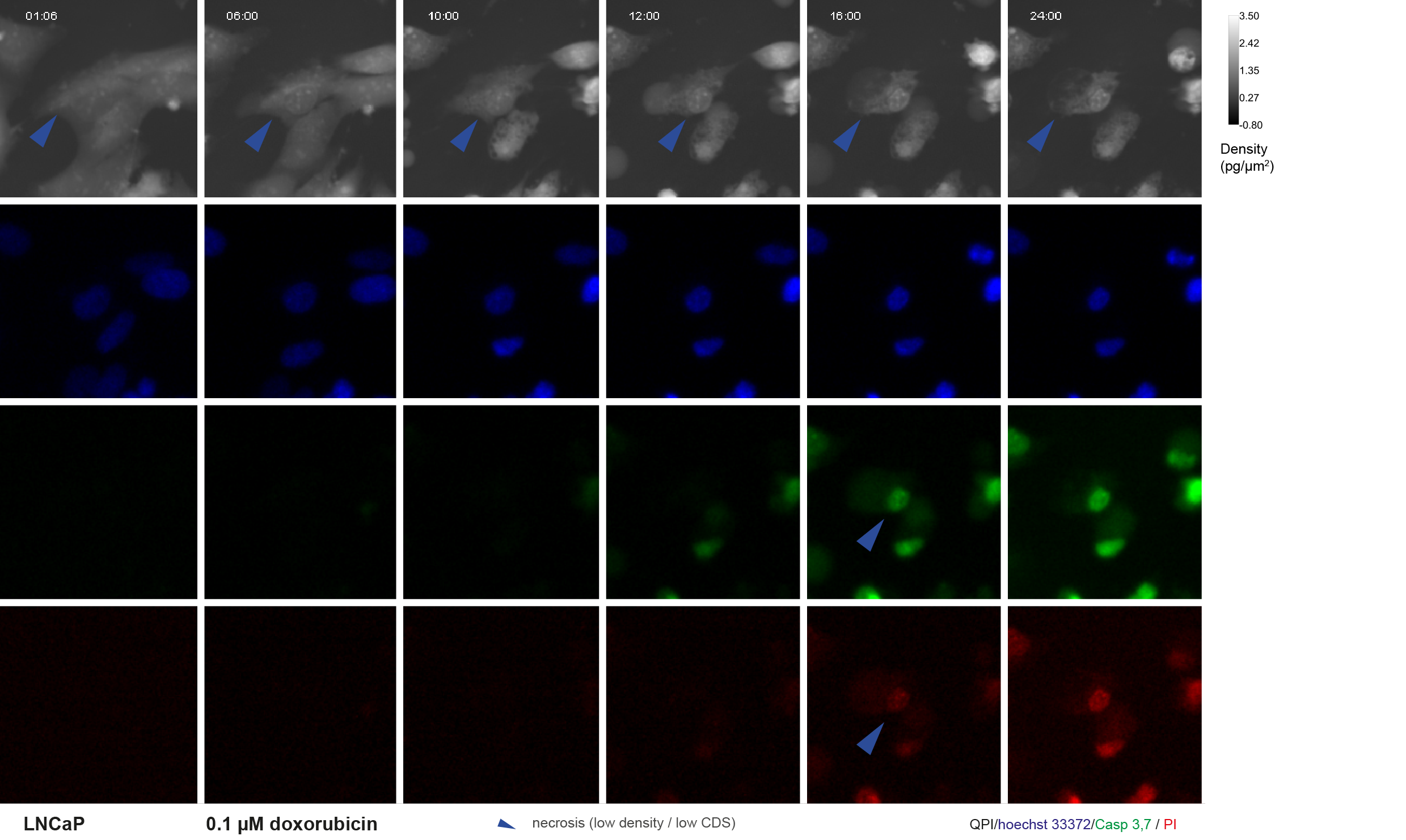

**Supplementary fig. 9:** **Critical timepoints for necrotic cell death - quantitative phase and fluorescence image (low cell density – low cell dynamic score)**. LNCaP cells exposed to 0.1 μM doxorubicin. Blue arrows indicate analysed necrotic cell (QPI image) and membrane rupture (fluorescence image). Notice simultaneous onset of PI and Casp 3, 7 signals. For quantitative time-lapse phase-related signals see supplementary fig. 8. 10x magnification. FOV size approx. 107 μm. QPI, quantitative phase image; PI, propidium iodide.

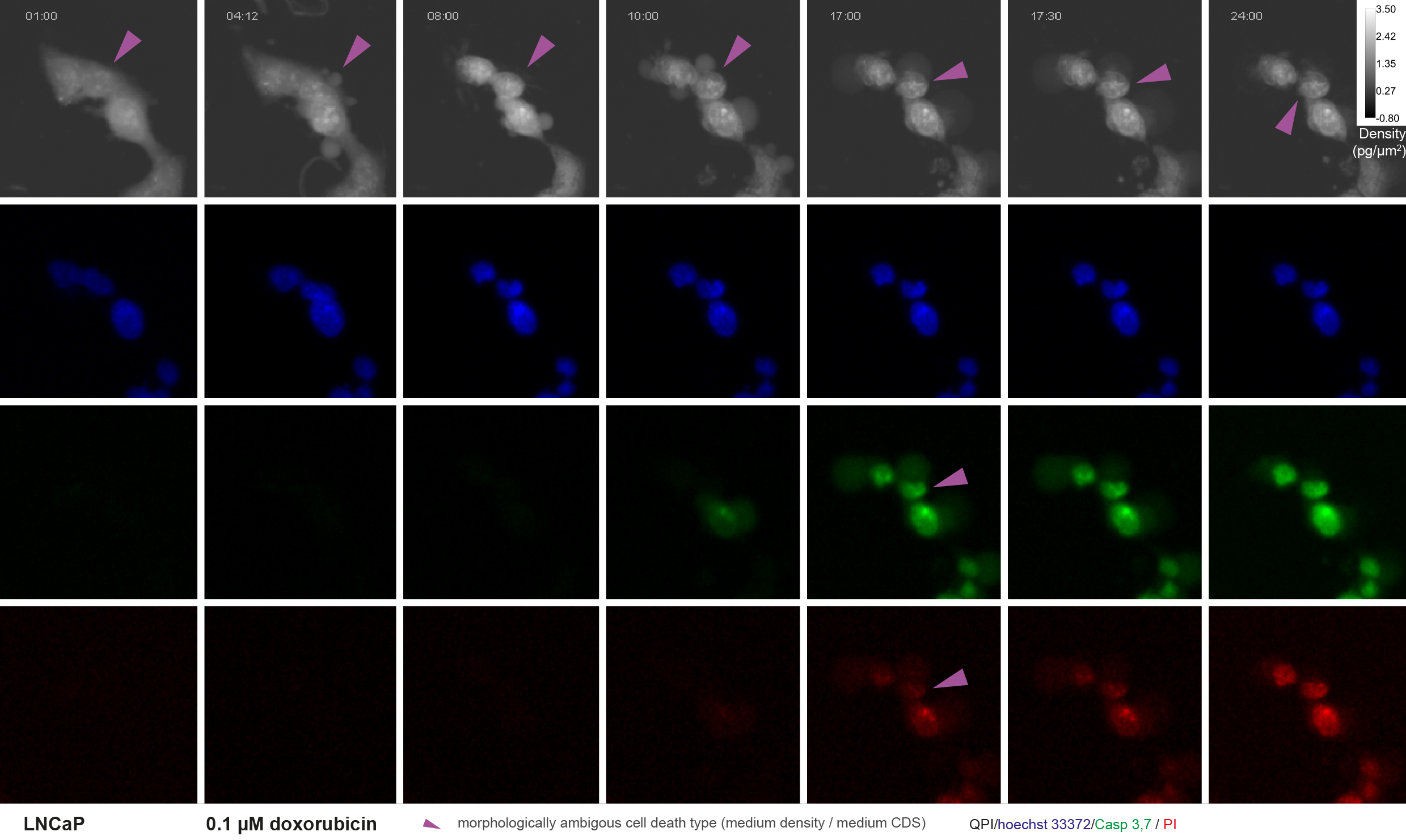

**Supplementary fig. 10: Critical timepoints for ambiguous cell death - quantitative phase and fluorescence image (medium cell density – medium cell dynamic score)**. PNT1A cells exposed to 0.1 μM doxorubicin. Violet arrows indicate analysed cell undergoing ambiguous death type (QPI image) and membrane rupture (fluorescence image). For quantitative time-lapse phase-related signals see supplementary fig. 8. 10x magnification. FOV size approx. 107 μm. QPI, quantitative phase image; PI, propidium iodide.

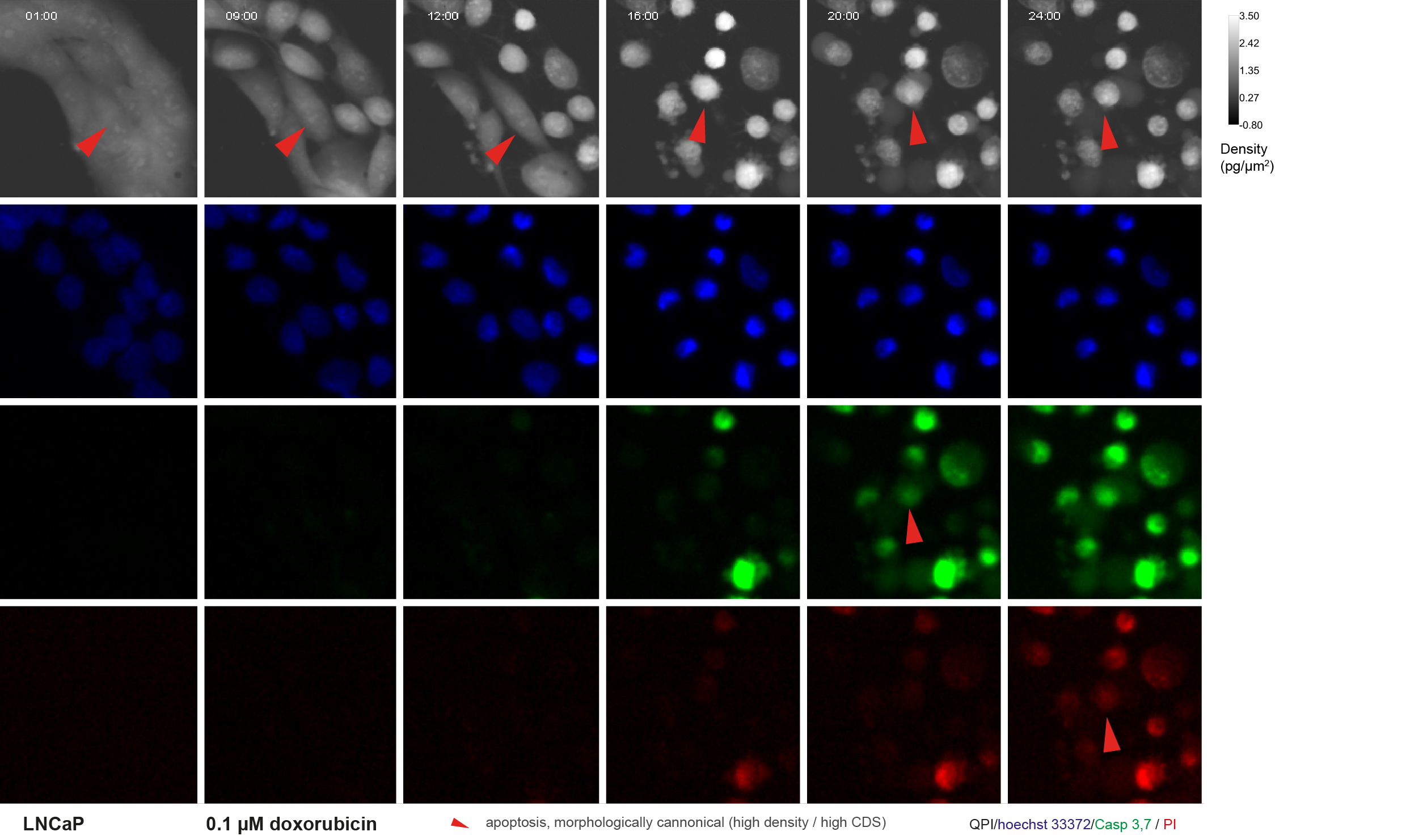

**Supplementary fig. 11: Critical timepoints for morphologically canonical apoptosis - quantitative phase and fluorescence image (high cell density – high cell dynamic score)**. LNCaP cells exposed to 0.1 μM doxorubicin. Red arrows indicate analysed cell undergoing cannonical apoptosis (QPI image) and membrane rupture (fluorescence image). Notice the onset of Casp 3,7 fluorescence before PI. For quantitative time-lapse phase-related signals see supplementary fig. 8. 10x magnification. FOV size approx. 107 μm. QPI, quantitative phase image; PI, propidium iodide.

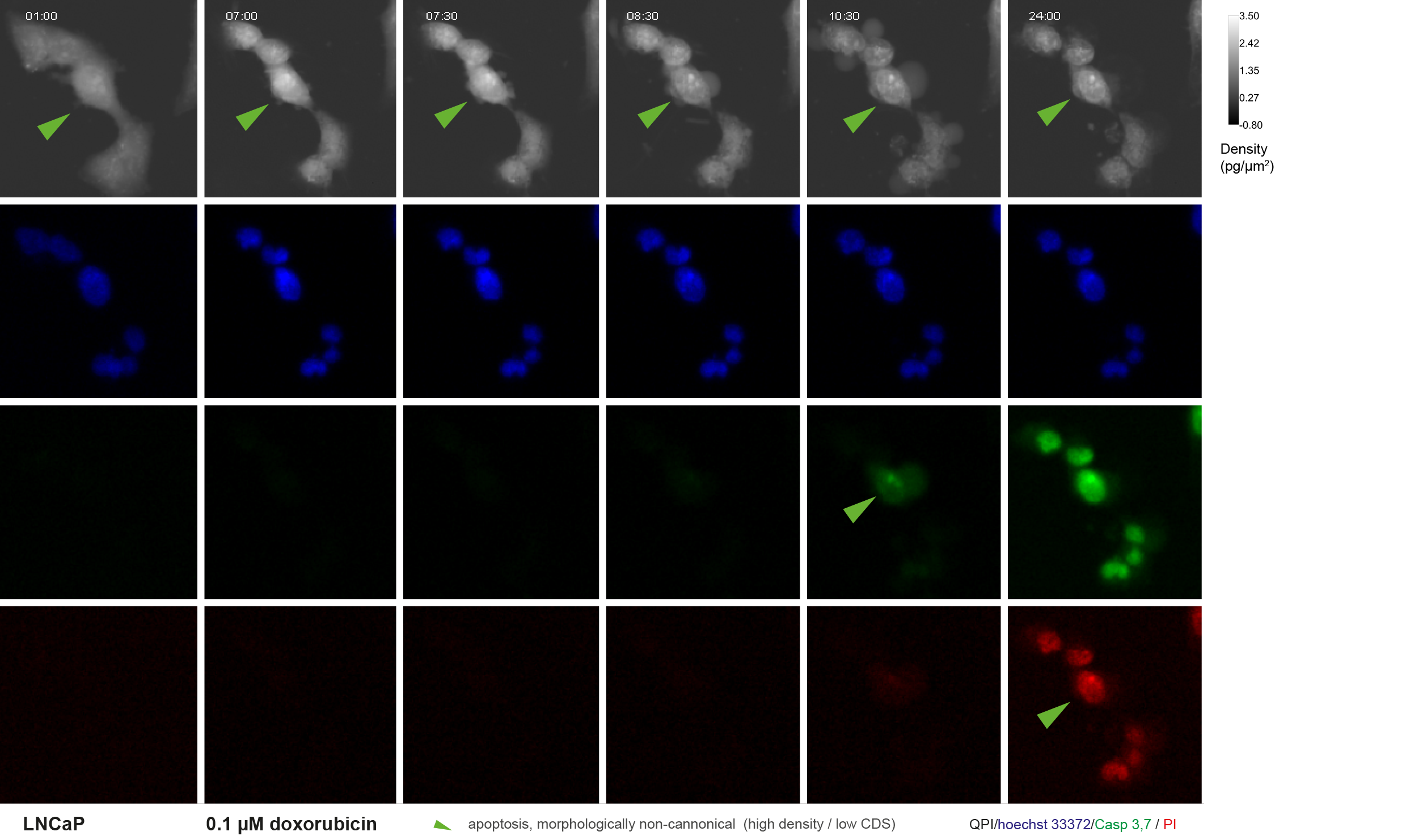

**Supplementary fig.12: Critical timepoints for morphologically non-canonical apoptosis - quantitative phase and fluorescence image**. LNCaP cells exposed to 0.1 μM doxorubicin. Red arrows indicate analysed cell undergoing cannonical apoptosis (QPI image) and membrane rupture (fluorescence image). For quantitative time-lapse phase-related signals see supplementary fig. 8. 10x magnification. FOV size approx. 107 μm. QPI, quantitative phase image; PI, propidium iodide.

**Supplementary table 1: Cell death type is characteristic by delayed onset of Caspase 3,7 and PI signal and by different nuclear Hoechst 33342 signal intensity.** Table of fluorescence signal intensities and Caspase 3,7 – PI signal onset delay, number of cells analyzed in manually and automatically annotated dataset of apoptotic and necrotic cells. Kruskal Wallis test used for p value calculation. PI, propidium iodide, CI, confidence interval.

| Cell line | treatment | classif. | death type | N cells | Hoechst 33342 avg. signal intensity | | Casp 3,7 – PI signal onset delay (frames) | |
| --- | --- | --- | --- | --- | --- | --- | --- | --- |
|  |  |  |  |  | median (95% CI) | p-val | median (95% CI) | p-val |
| DU-145 | doxorubicin | Manual | apoptosis | 86 | 240.19 (215.92 - 270.13) |  | 16.5 (7 - 111) |  |
|  |  | Manual | necrosis | 74 | 232 (216.44 - 248.87) | 0.5547 | 9 (5 - 15) | 0.0002 |
|  |  | Automatic | apoptosis | 72 | 245.45 (219.97 - 271.38) |  | 18 (8.5 - 129) |  |
|  |  | Automatic | necrosis | 88 | 229.87 (215.09 - 246.88) | 0.0154 | 9.5 (4 - 15) | 0.0001 |
|  |  |  | viable | 65 |  |  |  |  |
|  | black phosphorus | Manual | apoptosis | 39 | 769.5 (648.22 - 913.79) |  | 62 (14.5 - 139.5) |  |
|  |  | Manual | necrosis | 199 | 621.81 (535.37 - 701.19) | <0.0001 | 11 (1 - 19.75) | <0.0001 |
|  |  | Automatic | apoptosis | 32 | 815.99 (654.03 - 905.07) |  | 15 (4.5 - 55) |  |
|  |  | Automatic | necrosis | 206 | 625.25 (535.88 - 713.23) | <0.0001 | 12 (1 - 22) | 0.467 |
|  |  |  | viable | 35 |  |  |  |  |
|  | staurosporine | Manual | apoptosis | 105 | 253.88 (236.9 - 286.64) |  | 182 (117.75 - 250) |  |
|  |  | Manual | necrosis | 148 | 236.8 (221.41 - 257.76) | <0.0001 | 5 (1 - 11) | <0.0001 |
|  |  | Automatic | apoptosis | 15 | 285.6 (261.12 - 317.33) |  | 152 (98 - 246.25) |  |
|  |  | Automatic | necrosis | 238 | 243.91 (226.14 - 263.65) | 0.0015 | 11 (4 - 140) | 0.0074 |
|  |  |  | viable | 4 |  |  |  |  |
| LNCaP | doxorubicin | Manual | apoptosis | 61 | 276.75 (257.51 - 309.35) |  | 48 (14.75 - 133.25) |  |
|  |  | Manual | necrosis | 53 | 237.41 (223.97 - 267.13) | <0.0001 | 7 (1 - 16) | <0.0001 |
|  |  | Automatic | apoptosis | 29 | 290.22 (271.23 - 324.35) |  | 31 (14.75 - 117.5) |  |
|  |  | Automatic | necrosis | 85 | 245.07 (226.23 - 275.86) | <0.0001 | 12 (1 - 49.25) | 0.0077 |
|  |  |  | viable | 61 |  |  |  |  |
|  | black phosphorus | Manual | apoptosis | 42 | 847.83 (715.79 - 976.58) |  | 59 (12 - 160) |  |
|  |  | Manual | necrosis | 97 | 564.97 (498.39 - 667.04) | <0.0001 | 16 (1 - 60.25) | 0.0032 |
|  |  | Automatic | apoptosis | 31 | 798.04 (652.33 - 942.52) |  | 37 (6.5 - 109.5) |  |
|  |  | Automatic | necrosis | 108 | 577.93 (503.58 - 695.62) | <0.0001 | 21 (2.5 - 76) | 0.6102 |
|  |  |  | viable | 31 |  |  |  |  |
|  | staurosporine | Manual | apoptosis | 93 | 280.11 (255.52 - 309.28) |  | 75 (25.75 - 125.75) |  |
|  |  | Manual | necrosis | 107 | 258.87 (232.83 - 281.35) | 0.0001 | 11 (1 - 20) | <0.0001 |
|  |  | Automatic | apoptosis | 23 | 290.84 (254.05 - 333.54) |  | 37 (13.5 - 86.75) |  |
|  |  | Automatic | necrosis | 177 | 266.15 (245.21 - 289.56) | 0.0796 | 20 (8 - 76.25) | 0.9009 |
|  |  |  | viable | 36 |  |  |  |  |
| PNT1A | doxorubicin | Manual | apoptosis | 66 | 214.3 (195.92 - 254.59) |  | 49.5 (25 - 87) |  |
|  |  | Manual | necrosis | 167 | 206.7 (190.65 - 222.47) | 0.065 | 1 (1 - 5.75) | <0.0001 |
|  |  | Automatic | apoptosis | 30 | 244.13 (207.3 - 268.67) |  | 69.5 (29 - 90) |  |
|  |  | Automatic | necrosis | 203 | 206.29 (190.21 - 223.51) | 0.0001 | 1 (1 - 13) | <0.0001 |
|  |  |  | viable | 123 |  |  |  |  |
|  | black phosphorus | Manual | apoptosis | 132 | 748.23 (661.16 - 880.63) |  | 18 (14 - 63.5) |  |
|  |  | Manual | necrosis | 116 | 660.84 (598.24 - 715.6) | <0.0001 | 16 (1 - 54.5) | 0.022 |
|  |  | Automatic | apoptosis | 118 | 779.47 (675.22 - 900.14) |  | 17 (13 - 43) |  |
|  |  | Automatic | necrosis | 130 | 660.84 (598.16 - 707.82) | <0.0001 | 17.5 (9 - 62) | 0.9992 |
|  |  |  | viable | 31 |  |  |  |  |
|  | staurosporine | Manual | apoptosis | 22 | 263 (233 - 275.35) |  | 20.5 (1 - 95) |  |
|  |  | Manual | necrosis | 158 | 233.59 (219.37 - 252.52) | 0.0125 | 1 (1 - 20) | 0.0876 |
|  |  | Automatic | apoptosis | 22 | 260.99 (220.32 - 269.75) |  | 6 (1 - 64) |  |
|  |  | Automatic | necrosis | 158 | 234.36 (219.38 - 252.74) | 0.145 | 1 (1 - 28) | 0.6533 |
|  |  |  | viable | 4 |  |  |  |  |
| summary | all | Manual | apoptosis | 646 | 286.35 (245.91 - 658.55) |  | 55 (16 - 143) |  |
|  |  | Manual | necrosis | 1119 | 259.26 (224.19 - 563.72) | <0.0001 | 7 (1 - 17) | <0.0001 |
|  |  | Automatic | apoptosis | 118 | 278.65 (234.11 - 532.19) |  | 20.5 (11.5 - 94.5) |  |
|  |  | Automatic | necrosis | 130 | 278.3 (233.85 - 605.37) | 0.9326 | 11 (1 - 48) | <0.0001 |
|  |  |  | viable | 390 |  |  |  |  |
